# Antagonistic and Synergistic Roles of Tomato AFP3 Isoforms in Hormonal Regulation and Development

**DOI:** 10.1101/2025.06.18.660365

**Authors:** Ylenia Vittozzi, Louise Petri, Maurizio J. Chiurazzi, Purificación Lisón, Carmen Grech Hernández, Adity Majee, Naveen Shankar, Meike Burow, Stephan Wenkel

## Abstract

The ABA INSENSITIVE5 BINDING PROTEIN (AFP) family plays a critical role in abscisic acid (ABA) signaling through interaction with the transcription factor ABI5, impacting seed germination and stress responses. Here, we characterize tomato AFP3, which produces two isoforms: a full-length protein and a shorter microProtein (sAFP3) containing only the C-terminal domain. Functional analyses reveal contrasting roles of these isoforms in development; while *AFP3* overexpression accelerates shoot growth but impairs seed germination, both *afp3* loss-of-function and *sAFP3*-expressing mutants (*afp3-D*) exhibit stunted growth and developmental defects. Transcriptome profiling highlights that AFP3 and sAFP3 differentially regulate hormone-related pathways, including salicylic acid, gibberellic acid, and jasmonic acid metabolism. Proteomic interaction studies demonstrate that AFP3 and sAFP3 physically interact, sharing partners involved in hormone signaling. Hormone quantification confirms that AFP3 modulates multiple phytohormones, with elevated ABA in *afp3-D* mutants and increased gibberellic acid, jasmonic acid, and salicylic acid in both *afp3* and *afp3-D* mutant backgrounds. Moreover, AFP3 controls flower and fruit development, influencing yield and ripening. Together, these findings identify AFP3 as a key integrator of hormonal crosstalk that coordinates growth, development, and stress responses in tomato. The production of dual AFP3 isoforms through alternative transcription, combined with microProtein-mediated dominant-negative regulation, reveals a sophisticated mechanism for dynamically fine-tuning transcriptional networks. This versatile strategy underscores how plants—and potentially other organisms—achieve precise control over complex signaling pathways.

## INTRODUCTION

The use of high-throughput sequencing technologies and comprehensive transcriptomic analyses of eukaryotes has revealed that transcriptional regulation is more complex than previously thought. It is now well established that individual genes can produce multiple transcript isoforms through processes such as alternative splicing, alternative transcription initiation, and alternative polyadenylation^1^. These alternative isoforms may have different regulatory elements, splice variants, or untranslated regions, resulting in diverse functional outcomes^2^. Assigning biological functions to the various transcript isoforms that can be detected is a major challenge, and many of these isoforms may be byproducts with no biological relevance. Moreover, transcript isoforms that play a biological role may operate as RNA, protein, or both^3^.

Furthermore, the last decade has revealed additional transcripts that encode small proteins. These include small open reading frames (smORFs) located within non-coding RNAs, such as long non-coding RNAs (lncRNAs) or microRNAs (miRNAs) that can give rise to small proteins and peptides^4,5^. It is widely accepted that many of these small proteins play critical roles in growth and development and serve as disease biomarkers and therapeutic targets in human biology^6,7^.

The *ABA INSENSITIVE5 BINDING PROTEIN (AFP)* gene family is known to produce alternative transcripts^8^ and some AFP protein isoforms have been shown to act in a dominant negative fashion^9^. AFPs have long been known to interact with ABSCISIC ACID (ABA)-INSENSITIVE5 (ABI5)^10,11^. ABI5 is a basic leucine zipper transcription factor that regulates ABA signaling and controls germination^12^. AFPs are composed of three conserved domains called A-, B-, and C-domains. The A-domain contains an ethylene-responsive element binding factor-associated amphiphilic repression (EAR) motif, while the B-domain includes a classical bipartite nuclear localization signal (NLS). The C-terminal segment of AFPs is essential for interacting with ABI5 and other transcription factors (TFs), facilitating both homo- and heteromerization of AFPs^11,13^. Through the EAR domain, AFPs can recruit TOPLESS (TPL)/TOPLESS-RELATED (TPR) transcriptional co-repressor proteins that are central to plant development^14,15^. In contrast to jasmonic acid signalling where JAZ and NINJA proteins recruit the TPL/TPR complex, ABI5-BINDING PROTEINS (AFPs) recruit TPL/TPR proteins to _ABI5_^16–18^.

Despite initial studies indicating a pivotal role in germination and abiotic stresses such as drought and salinity, recent discoveries have highlighted the function of AFPs in other processes, including flowering regulation. The Arabidopsis AFP2 protein physically interacts with both the flowering time regulator CONSTANS (CO) and TOPLESS RELATED2 (TPR2) through the C- and A-domains, respectively. Thereby, AFP2, CO, and TPR2 form a trimeric complex that inhibits flowering by suppressing the expression of *FLOWERING LOCUS T*^13^. Most of our understanding of the molecular functions of the AFP gene family derives from studies in Arabidopsis. To extend this knowledge to fruit crop species, we initiated investigations in tomato.

Here, we investigated the functions of transcripts derived from the tomato *AFP3* gene. Initial database searches revealed an overrepresentation of ESTs covering the second exon of *AFP3*, prompting us to investigate the existence of alternative transcripts and their potential roles in tomato physiology and stress response. Specifically, we found that the tomato *AFP3* gene yields two major mRNA isoforms, the *AFP3* full-length transcript and s*AFP3* derived only from the second exon. Our findings reveal that *AFP3* and its splice variant *sAFP3* serve as key modulators of plant growth and development through distinct and interactive roles. Overexpression of AFP3 promotes shoot elongation, whereas overexpression of the sAFP3 microProtein causes developmental stunting and alters metabolic pathways. Despite these contrasting effects, the two isoforms can form heterodimeric complexes, suggesting coordinated involvement in hormonal signaling and transcriptional control. Their interplay manifests as both antagonistic and cooperative effects during development, highlighting the nuanced regulatory balance they maintain. Importantly, excessive AFP3 expression compromises plant vigour and productivity, underscoring the dose-sensitive nature of its function. Together, these findings position the AFP3-sAFP3 module as a critical node in the regulatory network that integrates growth, metabolism, and reproductive development.

## RESULTS

### Tomato AFP3 produces two mRNA isoforms

To examine the expression dynamics and functional significance of the tomato *AFP3* gene, we analyzed transcript variants and generated targeted gene deletions. Transcript characterization was performed using 5′-RACE PCR to identify mRNA isoforms produced from the *AFP3* locus. In *Arabidopsis thaliana*, previous studies demonstrated that *AFP1* mRNA is induced by exogenous flagellin^8^, whereas other AFP family transcripts are primarily responsive to abscisic acid (ABA)^11^. In wild-type tomato seedlings, *AFP3* 5′-RACE PCR products were not detected under untreated conditions, suggesting low or absent expression at this developmental stage (**Fig. 1A**). However, a 10-minute flagellin treatment induced the canonical *AFP3* transcript, while a four-hour ABA treatment led to the accumulation of both the canonical and an alternative transcript variant (**Fig. 1A**). To investigate AFP3 function, we generated CRISPR/Cas9-induced deletions at the *AFP3* locus (**Fig. 1B**). The tomato *AFP3* gene (*Solyc04g005380*) comprises two exons separated by an intron of approximately 3 kb (**Fig. 1C**). Exon 1 encodes the A-domain containing the EAR repression motif and the NLS-bearing B-domain, while exon 2 encodes the C-terminal domain. To determine the function of the *AFP3*-derived protein isoforms, we generated transgenic plants that overexpress the full-length AFP3 protein and the short C-domain containing microProtein isoform referred to as the sAFP3. In addition to overexpression, we used CRISPR-based genome engineering to create loss- and potential gain-of-function mutants. This was achieved by introducing a large deletion that started shortly after the translation start site and ended just before domain C (**Fig. 1B and C**). This deletion included domains A and B, as well as the large intron. Three loss-of-function mutations in the *AFP3* gene, referred to as *afp3-1*, *afp3-2*, and *afp3-3*, were obtained. These three mutants exhibit only minor differences from one another, but all lack the regions encoding domains A and B and were out of frame with domain C, thus rendering them null alleles (**Supplementary Fig. S1**). Additionally, we identified a putative gain-of-function allele, which we designated *afp3-D*. In comparison to *afp3-1*, *afp3-2*, and *afp3-3*, the reading frame in *afp3-D* remained unchanged after the genomic deletion. This resulted in a gene that had the same transcription/translation start as the original *AFP3* gene but exclusively produced the sAFP3 microProtein isoform.

**Figure 1.**
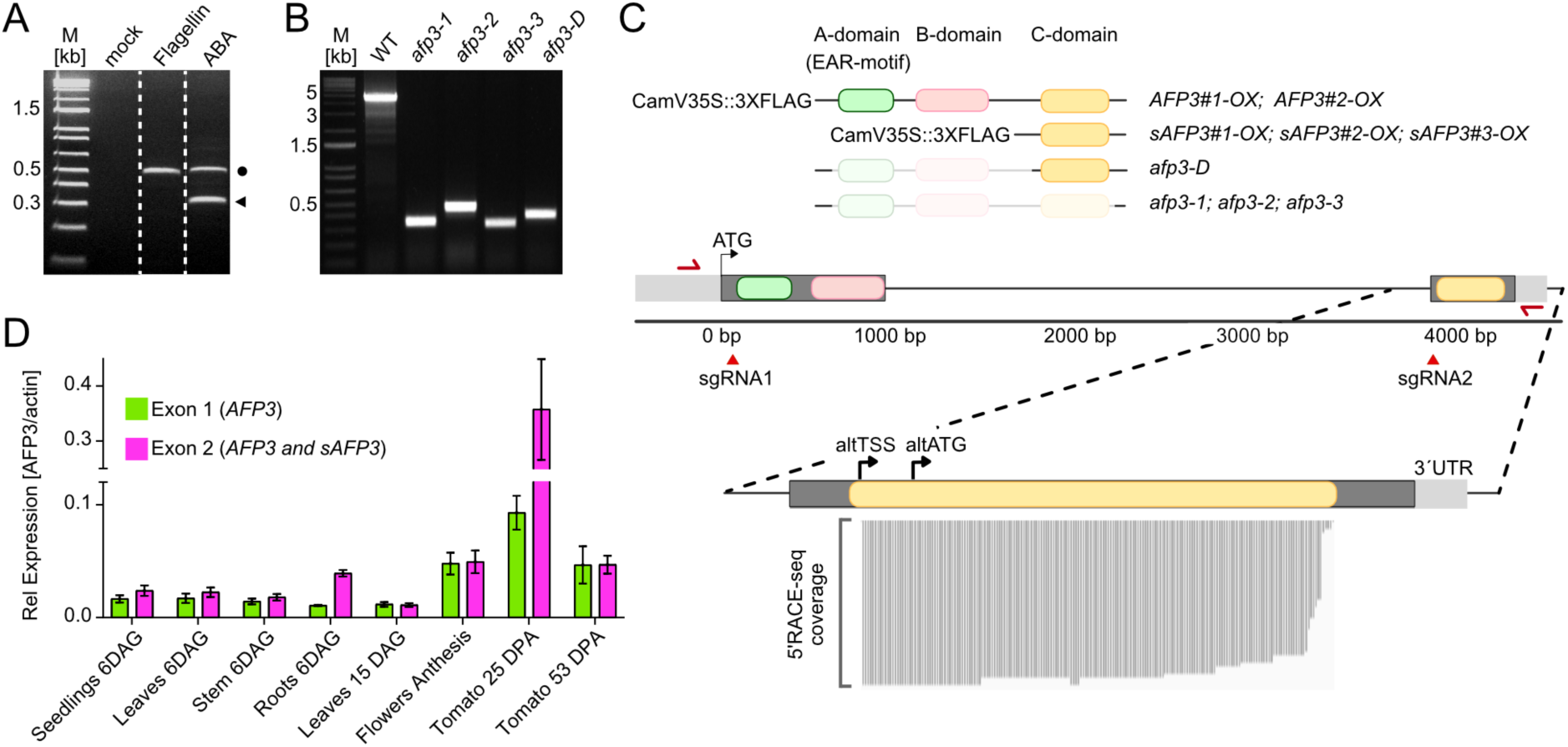
The tomato *AFP3* gene encodes two isoforms. **(A)** 5′RACE Nested PCR on WT seedlings subjected to different treatments (Flagelline 3,3 µM for 10 min, 100 µM ABA 4h, and mock, no treatment). The arrow highlights the identified alternative transcript, verified through Deep-sequencing of purified PCR product. **(B)** PCR gel image showing *AFP3* gene amplification in wild-type (WT) and CRISPR/Cas9 mutants, employing primers in panel A (red arrows). All mutants exhibit homozygous deletions. **(C)** <u>Upper panel:</u> Scaled schematic depiction of the tomato *AFP3* (*Solyc04g005380)* gene structure. In green, pink and yellow are respectively depicted the Ear-motif, B-domain, and C-domain in the two exons, conserved domains of AFP gene family. Above, predicted protein structures of the CamV35SxFLAG fused proteins (*AFP3-OX* and *sAFP3*), and CRISPR/Cas9 mediated mutants, including an *in-frame deletion* of the EAR and B-domain (*afp3-D*) and the knockouts due to deletion and additional frameshifts (*afp3-1, afp3-2, afp3-3*). Red triangles mark Cas9 target sites, and red arrows indicate the oligos used for PCR genotyping of CRISPR mutants. In magenta and dark green are the regions detected by RT-qPCR represented in panel D. <u>Lower panel:</u> Schematic representation of alternative transcript (*sAFP3*) within the magnified second exon of *AFP3* gene. Red arrows indicate the reverse primers for the 5′RACE and Nested PCR. Black arrows show the alternative Transcription Start Site (altTSS) and alternative (altATG) identified by Deep sequencing. **(D)** Tissue specific qRT-qPCR showing absolute expression of *AFP3* gene regions in WT plants. Magenta and dark green represent specifically exon 1 (*AFP3*) and exon 2 (*AFP3+sAFP3*), see panel C.

To validate the production of the alternative transcript and obtain information on the transcription start site, we performed 5’RACE-seq. After enriching the alternative transcript by RACE-PCR, we generated a sequencing library and used Illumina short-read sequencing to determine the exact sequence start of the alternative transcript. Deep sequencing revealed that all identified transcripts linked to the 5′RACE adaptor started at the same position, which included an alternative translation start 12bp downstream of the transcription start site (**Fig. 1C** and **Suppl. Fig. S2**).

Analysis of the mRNA expression in different developmental stages and tissues revealed that *AFP3* is expressed throughout plant development. The highest expression levels were observed in green fruits (25 days post anthesis, DPA), which is in agreement with the gene expression atlas (Tomato Expression Atlas, Sol Genomics), which reports higher *AFP3* expression in the early stages of immature fruits after anthesis. Expression of *sAFP3* does not differ from that of *AFP3*, except in six-day-old roots and 25 DPA green tomato fruits, where its expression was higher (**Fig. 1D**). These findings indicate that the *sAFP3* transcript is dynamically expressed during development and stress.

### AFP3 and sAFP3 show contrasting developmental defects

Short proteoforms, such as microProteins derived from alternative transcripts, can regulate transcription factor activity by forming non-productive protein complexes^19–21^. These proteoforms can, for example, bind the full-length isoform, forming non-productive protein complexes. To better understand the function of the tomato *AFP3* gene and its smaller isoform, we examined the growth behavior of both mutant plants and transgenic plants overexpressing either the full-length AFP3 protein or the truncated sAFP3 microProtein isoform. Phenotypic scoring was performed using T3 homozygous knockout plants (*afp3-1*) and the potential gain-of-function mutants (*afp3-D*). Additionally, T2 transgenic overexpression plants (*AFP3-OX* and *sAFP3-OX*) were used, derived from selected T0 lines based on their gene and protein expression levels. (**Suppl. Fig. S3A**). Compared to wild-type plants, we observed reduced growth and stunted development in *afp3-D* and *afp3-1* plants, as well as in the three independent *SAFP3-OX*lines. Conversely, the two independent transgenic lines, *AFP3#1OX* and *AFP3#2-OX*, exhibited accelerated shoot growth, resulting in taller plants (**Fig. 2A,B**). However, transgenic plants expressing the *AFP3* isoform at high levels (*AFP3-OX*) produced a high percentage of non-germinating seeds indicative of a negative role of AFP3 in seed germination. Genetic testing of the transgenic offspring showed that all plants taller than the wild type were heterozygotes. In contrast, a small number of stunted and pale plants were identified as homozygous overexpressors (**Suppl. Fig. S3B**). We performed phenotyping experiments with heterozygote plants because seeds from homozyogous plants exhibited very poor germination rates. Since the *AFP3-OX* transgene induces a dominant phenotype, we focused on phenotyping heterozygous plants due to their stable growth characteristics. The strongest growth suppression was observed in plants expressing the sAFP3 isoform, either transgenically or endogenously in the *afp3-D* mutants generated via genome engineering (**Fig. 2A, B**). Based on the phenotype observed 33 days after germination (DAG) and onwards, the genome-engineered *afp3-D* plants were indistinguishable from the transgenic *sAFP3-OX* plants. These findings reveal that distinct AFP3 isoforms have opposing effects on plant development—promoting shoot growth or suppressing it—highlighting AFP3 as a key regulatory node with isoform-specific functions.

**Figure 2.**
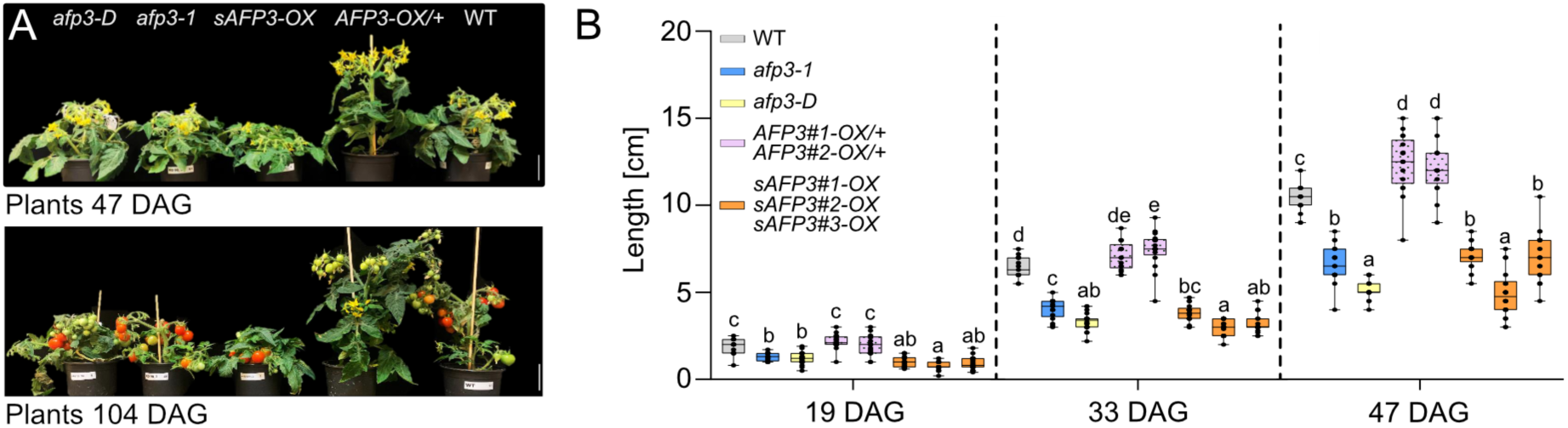
The AFP3 and sAFP3 proteoforms have opposing functions in growth regulation. **(A)** Picture of representative transgenic and mutant plants *afp3-D, afp3-1, sAFP3-OX*, *AFP3-OX/+* and WT during phenotypic characterization at 47 and 104 Days After Germination (DAG) (bar=5 cm). **(B)** Shoot Length measures represented as a Time course of 19, 33, and 47 DAG. The analysis includes WT (grey), homozygous knockout *afp3-1* (blue), homozygous *afp3-D (yellow), two* heterozygous overexpression lines of the full-length protein *AFP3#1-OX/+ and AFP3#2-OX/+* (pink)*, three homozygous overexpression lines of microProtein sAFP3#1-OX, sAFP3#2-OX, sAFP3#3-OX* (orange). One-way ANOVA with post hoc Tukey Test multiple comparisons for individual time points is made in R studio. Confidence level 0.95, significance alpha=0.05 (P < 0.05). Graphical representation of box plots in GraphPad Prism represents individual data points in addition to mean + SEM (n= 15 control plants, n=15 to 21 mutant plants).

Other growth phenotypes were also evaluated in the respective transgenic and mutant plants, including the number and length of axillary shoots, leaf morphology, and leaf primary axis length (**Supp. Fig. S4**). As previously stated, heterozygote plants overexpressing *AFP3* exhibited increased shoot growth, while plants overexpressing the *sAFP3* isoform exhibited reduced growth. This growth pattern was consistent across independently transformed plants (**Suppl. Fig. S4A**). Heterozygous plants overexpressing *AFP3* exhibited a significant decrease in axillary shoot formation at 47 DAG, characterized by both a reduced number and shorter length of the shoots (**Suppl. Fig. S4B, C, E**). Additionally, the primary axis length of leaves was reduced in all mutants and transgenic plants we generated. As a reference, we measured the size of the fourth composite leaf of each plant (**Suppl. Fig. S4D, F**). Finally, we also noted changes in leaf morphology. Mild defects in leaflet positioning were observed in *afp3* mutants and heterozygous transgenic plants overexpressing *AFP3*, whereas *afp3-D* mutants and transgenic plants overexpressing *sAFP3* exhibited distinct leaflet twisting phenotypes (**Suppl. Fig. S4D**). These findings further support the dominant effect of sAFP3-induced phenotypic alterations.

### Transcriptional network regulated by AFP3 and sAFP3

Ectopic expression of AFP3 induces growth, while sAFP3 acts to suppress growth. To determine whether the observed phenotypic changes are associated with differences in gene expression, we conducted a genome-wide transcriptome analysis. This analysis compared wild-type plants with *afp3* and *afp3-D* mutants, as well as *AFP3-OX* and *sAFP3-OX* transgenic lines. RNA of six-day-old seedlings grown under long-day conditions was isolated and sequenced.

In total, we found 3873 differentially expressed genes across all genotypes (**Fig. 3A**). Initial principal component analysis (PCA) revealed five distinct clusters corresponding to the different genetic backgrounds **(Fig. 3B**). While four of the genotypes, *afp3-1*, *afp3-D*, *sAFP3-OX* and WT, grouped closely together, indicating relatively similar global expression profiles, the *AFP3-OX* samples formed a clearly separated cluster, suggesting a markedly different transcriptomic signature. One replicate of *afp3-D* was dissimilar to the other biological replicates and was therefore removed from further analysis. The largest number of DEGs was observed in the transgenic plants overexpressing *AFP3* (**Fig. 3C**). Gene ontology (GO) analysis of genes upregulated in *AFP3-OX* transgenic plants revealed a significant enrichment of genes associated with abaxial cell fate specification, followed by those involved in salicylic acid biosynthesis and signaling (**Supplementary Dataset 1**). Since AFP3 contains an EAR motif, it is likely to function as a transcriptional repressor. To explore this further, we examined the GO categories of genes downregulated in the respective transgenic lines. This analysis revealed a significant enrichment of genes associated with osmotic and salt stress responses. In contrast, overexpression of the sAFP3 microProtein led to the deregulation of various metabolic pathways, including upregulation of genes involved in auxin biosynthesis and downregulation of those related to jasmonic acid metabolism (**Supplementary Dataset 1 and 2**).

**Figure 3.**
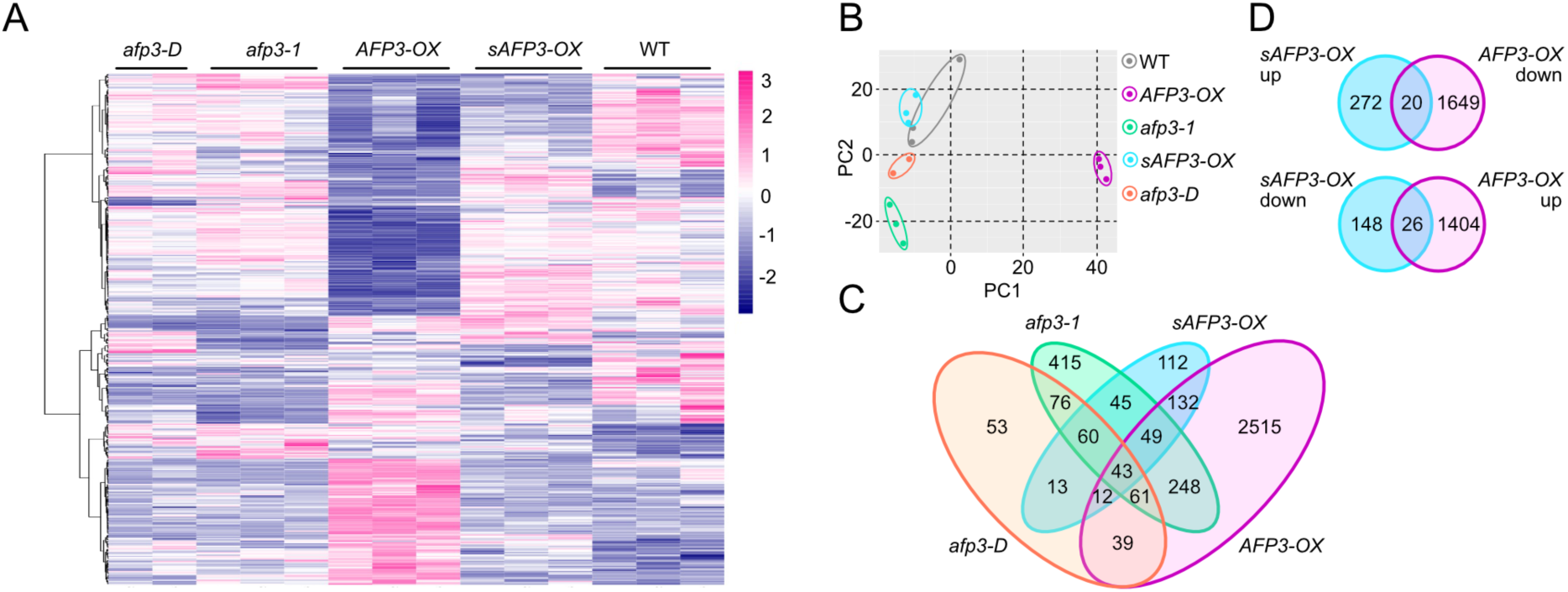
**Characterization of the transcriptional network regulated by AFP3 and sAFP3**. **A)** Heatmap of Differentially Expressed Genes (DEG)s in the different genotypes included in the transcriptomic data. Color scale represents difference between highly expressed genes (magenta), and lowly expressed genes (blue). **(B)** Principal component analysis (PCA) of the gene expression data shows similarities between biological replicates for each genotype, except for *afp3-D* where one biological replicate was excluded from the analysis as it was found to be an outlier after sequencing. **(C)** Venn-diagram showing all DEGs in the genotypes included in the RNA-seq compared to the WT. **(D)** Venn Diagrams showing the common genes that are both up-regulated and down-regulated in *AFP3-OX* compared with *sAFP3-OX*.

Given that *sAFP3* overexpression produced a phenotype opposite to that of *AFP3* overexpression, we further analyzed genes that were differentially regulated in opposite directions by *AFP3* and *sAFP3* overexpression. This analysis revealed 20 genes that were upregulated by sAFP3 and downregulated by AFP3, and 26 genes that were downregulated by sAFP3 and upregulated by AFP3 (**Fig. 3D**). Upon closer inspection of the 20 genes that were regulated in opposite directions, it was found that most of them encoded enzymes related to amino acid metabolism (**Table 1**). Importantly, the 26 genes that were downregulated by AFP3 but upregulated by sAFP3 showed a significant overrepresentation of transcription factors involved in developmental regulation, as well as enzymes associated with gibberellin biosynthesis (**Table 1**). Since AFP3 is not expected to bind DNA directly but likely functions as a co-repressor within a transcription factor complex, it is plausible that these 26 downregulated genes represent direct targets of the AFP3-containing repressor complex.

**Table 1:** Gene showing opposite expression in AFP3/sAFP3-overexpression.

| AFP3 | sAFP3 | Gene ID | Annotation |
| --- | --- | --- | --- |
| up | down | Solyc12g098540 | Apyrase |
| up | down | Solyc08g080570 | UDP-glucose 4-epimerase |
| up | down | Solyc10g084120 | plasma membrane intrinsic protein 2.5 |
| up | down | Solyc08g068600 | Aromatic amino acid decarboxylase 1B |
| up | down | Solyc08g077810 | Cationic amino acid transporter, putative |
| up | down | Solyc04g076630 | Rhamnogalacturonate lyase family protein |
| up | down | Solyc09g090900 | Aconitate hydratase |
| up | down | Solyc10g050323 | uncharacterized |
| up | down | Solyc06g008620 | tolB protein-like protein |
| up | down | Solyc03g093550 | Ethylene-responsive transcription factor |
| up | down | Solyc05g047580 | Terpene cyclase/mutase family member |
| up | down | Solyc07g009100 | defensin-like protein |
| up | down | Solyc04g078110 | serine protease SBT1 |
| up | down | Solyc04g076460 | Leucine-rich repeat receptor-like protein kinase family protein |
| up | down | Solyc10g084240 | Peroxidase |
| up | down | Solyc03g121540 | beta-galactosidase 3 |
| up | down | Solyc10g007960 | Allene oxide synthase |
| up | down | Solyc12g027540 | Retrovirus-related Pol polyprotein from transposon TNT 1-94 |
| up | down | Solyc04g078290 | Cytochrome P450 |
| up | down | Solyc07g017610 | Lysine-ketoglutarate reductase/saccharopine dehydrogenase |
| down | up | Solyc05g012360 | alpha/beta-Hydrolases superfamily protein |
| down | up | Solyc12g088250 | Serine carboxypeptidase, putative |
| down | up | Solyc06g076400 | Protein phosphatase 2C |
| down | up | Solyc11g011980 | Transducin/WD40 repeat-like superfamily protein |
| down | up | Solyc09g007460 | POLAR LOCALIZATION DURING ASYMMETRIC DIVISION |
| down | up | Solyc08g006990 | aluminum activated malate transporter family protein |
| down | up | Solyc04g074920 | Photosystem I assembly protein Ycf3 |
| down | up | Solyc06g035530 | gibberellin 20-oxidase-2 |
| down | up | Solyc06g051940 | Protein phosphatase 2c, putative |
| down | up | Solyc09g091600 | Tetratricopeptide repeat-like superfamily protein |
| down | up | Solyc10g081810 | MD-2-related lipid recognition domain-containing protein |
| down | up | Solyc07g150103 | uncharacterized |
| down | up | Solyc03g096260 | Protein yippee-like |
| down | up | Solyc10g008270 | bHLH transcription factor 094 |
| down | up | Solyc12g039080 | Leucine-rich repeat protein kinase family protein |
| down | up | Solyc04g077450 | DNA binding protein |
| down | up | Solyc04g005800 | Homeobox leucine zipper family protein |
| down | up | Solyc11g065520 | Leucine-rich repeat protein kinase family protein |
| down | up | Solyc07g039410 | EDNR2GH3 protein |
| down | up | Solyc07g056570 | NCED3-like\|nced |
| down | up | Solyc10g083380 | BZIP transcription factor family protein |
| down | up | Solyc01g111970 | L-ascorbate oxidase-like protein |
| down | up | Solyc03g094000 | Light induced like protein |
| down | up | Solyc08g062290 | Light-independent protochlorophyllide reductase subunit B |
| down | up | Solyc06g062460 | bHLH transcription factor136 |
| down | up | Solyc09g091390 | GATA transcription factor 18 |

### sAFP3 interacts with AFP3 to control its activity

Given the opposing roles of AFP3 and sAFP3 in developmental regulation, we next examined their subcellular localization to gain further insight into their distinct cellular functions. To assess their localization, we transiently delivered eGFP-tagged versions of AFP3 and sAFP3, driven by the *CaMV 35S* promoter, into *Nicotiana benthamiana* leaves, using an eGFP empty vector as a control. Consistent with predictions from the bioinformatic tool LOCALIZER, AFP3 was found to localize predominantly to the nucleus, likely due to the presence of a nuclear localization signal (NLS) within its B domain. In contrast, transient expression of sAFP3, which lacks the B domain, resulted in both nuclear and cytoplasmic localization (**Fig. 4A**). To investigate whether AFP3 and sAFP3 co-localize within the cell, we performed co-localization experiments by co-expressing eGFP-tagged AFP3 with either mCherry-tagged AFP3 or mCherry-tagged sAFP3. AFP3 was found to localize exclusively to the nucleus and co-localized with itself when co-expressed with mCherry-AFP3. When co-expressed with mCherry-sAFP3, nuclear co-localization was also observed; however, a portion of sAFP3 remained in the cytoplasm, while AFP3 was never detected outside the nucleus (**Fig. 4B**). Notably, sAFP3 was unable to redirect AFP3 to the cytoplasm, suggesting that their interaction does not affect AFP3’s nuclear localization. Conversely, the nuclear presence of sAFP3 in co-expression conditions may result from its interaction with AFP3, implying that AFP3 could help recruit sAFP3 into the nucleus. These distinct yet partially overlapping localization patterns may underlie their antagonistic roles in development, with AFP3 acting primarily in nuclear repression and sAFP3 potentially modulating this activity through partial nuclear co-localization and other cytoplasmic functions.

**Figure 4.**
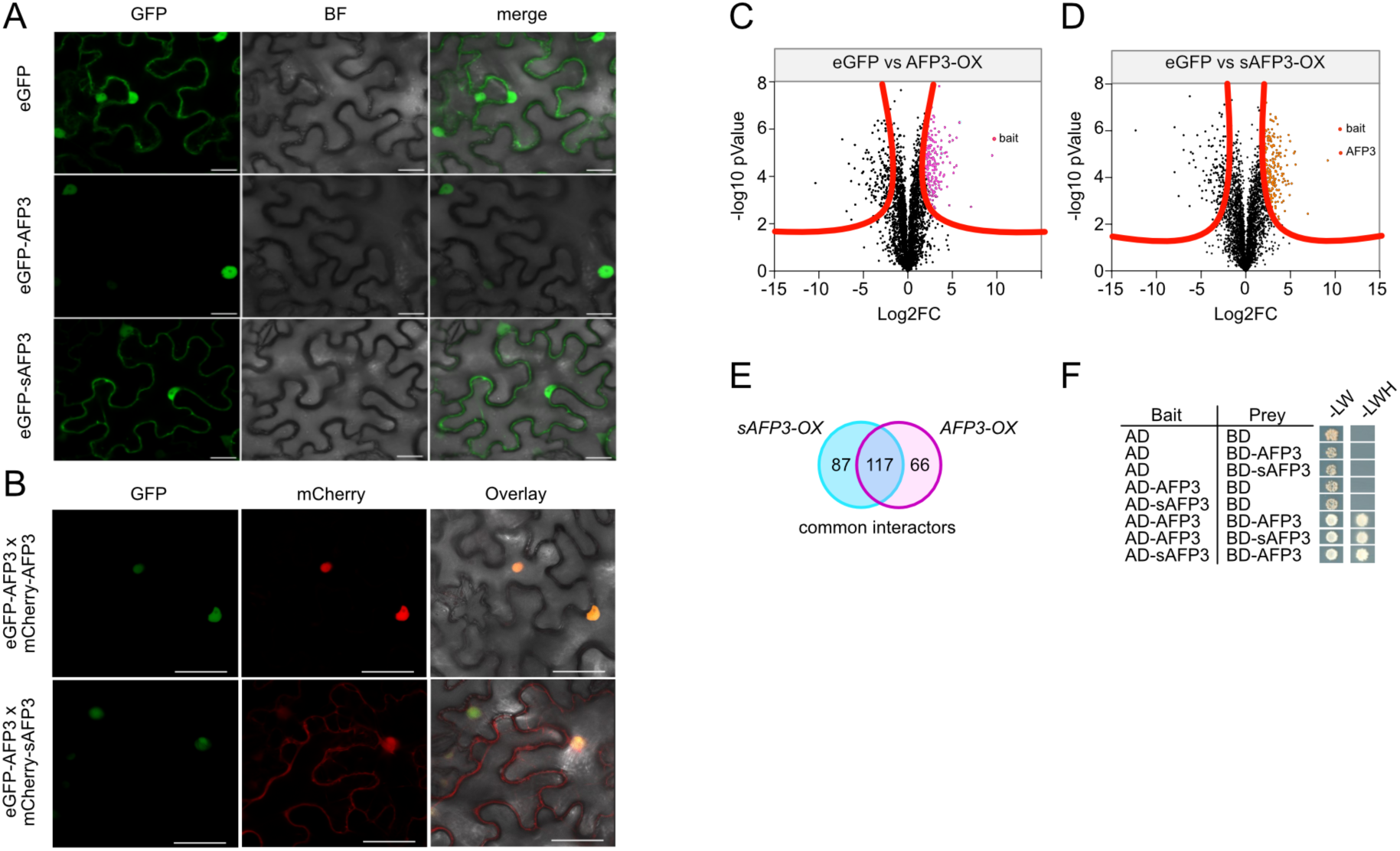
Identification of AFP3 and sAFP3 interacting proteins. **(A)** Subcellular localization of eGFP-AFP3 and eGFP-sAFP3 in *N. benthamiana* leaves. Panel show AFP3 localized to the nucleus, while sAFP3 was found in both cytoplasm and nucleus. Scale bar= 15 µm. **(B)** Co-localization of AFP3 and sAFP3 in transiently transformed tobacco leaves. Upper panel shows co-localization of eGFP-AFP3 and mCherry-AFP3 in nuclei; lower panel shows co-localization of eGFP-AFP3 with mCherry-sAFP3 in the nucleus. Note some mCherry-sAFP3 signal is also present in the cytoplasm. **(C,D)** Volcano plots show mass spectrometry analysis of protein enrichment after immunoprecipitation of Flag-tagged proteins in AFP3-OX and sAFP3-OX transgenic plants compared to eGFP-expressing control plants. Enriched proteins were identified by FDR-corrected test (FDR=0.01) and distributed on the graph according to the Log2 fold change of the average spectral counts and -Log10 (pValue). The magenta (AFP3-OX) and orange (sAFP3-OX) data points represent proteins that are considered significant with a log2 fold-change FC>2 and p-value<0.01. The detected baits and the full-length protein AFP3 in *sAFP3-OX* plants are shown in red. **(E)** Venn Diagram of the significantly detected proteins in both genotypes. *sAFP3-OX* and *AFP3-OX* showed 117 joint interactors, while respectively interacting with 87 and 66 different proteins. **(F)** Directed yeast-two-hybrid test showing that AFP3 can form homodimers in yeast and also heterdimerizes with sAFP3.

The observed antagonistic phenotypes between the transgenic and mutant plants suggest that sAFP3 counteracts AFP3 function, potentially by interfering with its activity. Their nuclear co-localization supports this possibility, placing sAFP3 in the same subcellular compartment as AFP3, where it may modulate its function through direct or indirect interaction. Considering that both protein isoforms contain domain C, they may form dimeric complexes via this region. To test this hypothesis and to identify other interacting proteins, we performed individual affinity purification mass spectrometry (AP-MS) experiments. To ensure comparability between datasets, we used the same transgenic plants and growth stages employed in the transcriptome studies. As a control, we conducted AP-MS experiments with wild-type and transgenic plants that overexpressed the FLAG-eGFP protein (*35S::FLAG-eGFP*). A total of 183 significantly enriched proteins were identified as interacting with AFP3 (**Fig. 4C**), while 204 proteins were found to interact with sAFP3 (**Fig. 4D**). The comparative analysis of both datasets revealed 117 proteins that were enriched with both baits (**Fig. 4E**). Surprisingly, these common interactors are predominantly proteins with functions in the chloroplast (**Supplementary Dataset 2**). This may suggest that these interactors are false positives or that AFP3/sAFP3 have a role in chloroplast biology. Among the 117 interactors, ARS2 and Gibberellin-regulated protein 6 function in ABA and gibberellic acid signaling, respectively, and a LOB domain transcription factor was also identified (**Supplementary Dataset 3**). These findings further support a recurring role for AFP3/sAFP3 in hormone-related processes, particularly in signaling pathways. This notion is reinforced by the observation that homozygous *AFP3-OX* transgenic plants exhibit impaired germination and various pleiotropic growth defects. Moreover, the identification of a LOB domain transcription factor as a common interactor suggests that AFP3 may function in transcriptional regulation as part of a LOB-domain protein complex. Furthermore, AFP3 protein was detected in the sAFP3 immunoprecipitations, which supports the formation of AFP3/sAFP3 heterodimeric complexes *in planta*. This may help explain the antagonistic effects observed in the developmental phenotypes associated with *AFP3* and *sAFP3* overexpression. To validate that AFP3 and sAFP3 can form heterodimers, we conducted directed yeast-two-hybrid studies. In these experiments, both AFP3 and sAFP3 were fused with either the Gal4 activation domain (AD) or the Gal4 DNA-binding domain (BD). We observed yeast growth on a selective medium lacking histidine, indicative of an interaction when AFP3 was present as both AD- and BD-fusion, which supports the hypothesis that AFP3 can dimerize. Additionally, we observed interactions of sAFP3 with AFP3, which supports the hypothesis that a direct physical interaction between AFP3 and sAFP3 is mediated through the domain C (**Fig. 4F**).

### AFP3 Modulates Multiple Phytohormones to Regulate Growth

Transcriptomic and AP-MS proteomics analyses implicated AFP3 in the regulation of hormone biology, particularly in the biosynthesis and signaling of abscisic acid (ABA), gibberellic acid (GA), and salicylic acid (SA). To validate AFP3’s role in hormone homeostasis and to determine whether the phenotypic alterations observed in *afp3* mutants could be attributed to changes in hormone profiles, we employed a comprehensive hormonomics approach. This involved quantifying the levels of key phytohormones in wild-type, *afp3* loss-of-function, and *afp3-D* mutant plants. Our analysis revealed significant perturbations in four major plant hormones. ABA levels were markedly elevated in *afp3-D* mutants, whereas no significant difference was detected between wild-type and *afp3* null mutants (**Fig. 5A**). In contrast, GA, jasmonic acid (JA), and SA levels were consistently increased in both *afp3* and *afp3-D* mutants relative to wild type (**Fig. 5B-D**). Given that *afp3-D* is a dominant-negative allele that impairs AFP3 function, it likely exerts its effects in part through interference with other AFP-like proteins. The distinct ABA phenotype—unaltered in *afp3* nulls but elevated in *afp3-D* mutants—suggests that ABA regulation involves complex mechanisms potentially extending beyond AFP3 function alone, possibly implicating functional redundancy within the AFP family. Importantly, the elevated levels of GA, JA, and SA observed in both mutant lines underscore AFP3’s role as an integrative regulator of multiple hormone biosynthetic and signaling pathways. This multilayered hormonal modulation likely underpins AFP3’s specific influence on tomato growth and developmental processes. Collectively, these findings position AFP3 as a critical node coordinating hormone crosstalk essential for proper developmental regulation.

**Figure 5.**
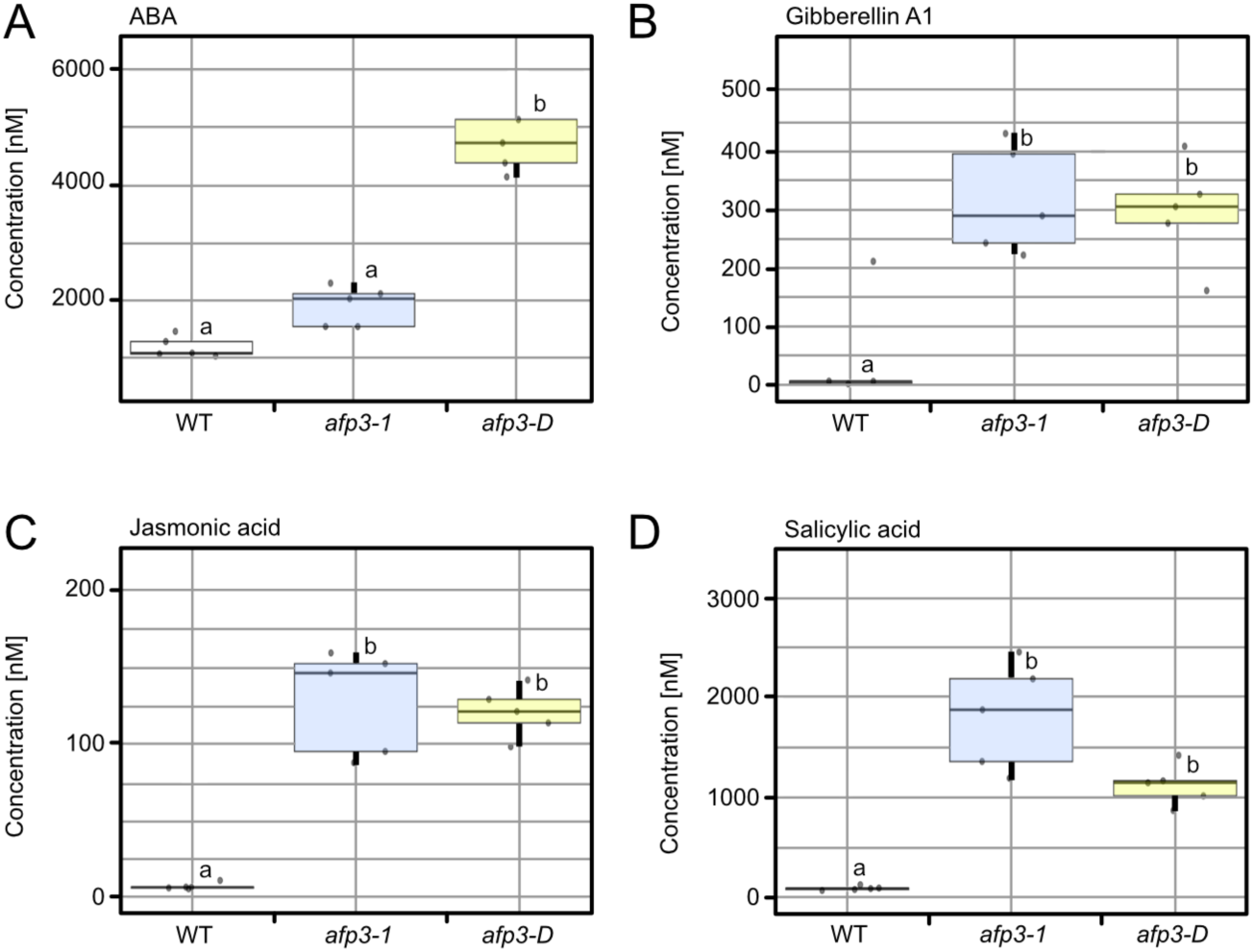
**Hormone profiling in 30-day-old tomato leaves**. Phytohormone levels were quantified in leaves of 30-day-old WT, *afp3-1*, and *afp3-D* plants using targeted hormone profiling via LC-QqQ-MS/MS. Shown are the concentrations of (**A**) abscisic acid (ABA), (**B**) gibberellin A_1_ (GA_1_), (**C**) jasmonic acid (JA), and (**D**) salicylic acid (SA). Statistical analysis was carried out in R using one-way ANOVA followed by Tukey’s HSD post hoc test for multiple comparisons. Different letters denote statistically significant differences (p < 0.05).

### Distinct Roles of AFP3 and sAFP3 in Flower and Fruit Development

Given the dynamic expression of *AFP3* and *sAFP3* in reproductive organs (**Fig. 1E**), we conducted a detailed phenotypic analysis at later developmental stages, revealing abnormalities in flower and fruit development. A positive effect on flower size was observed in transgenic *AFP3-OX* plants, which exhibited enlarged flowers at the anthesis stage compared to wild-type and *afp3* mutant plants (**Fig. 6A**). The fruit set measured in the first three influorescences varied among the genotypes analyzed. Notably, *AFP3-OX* heterozygous plants showed a significantly higher fruit set relative to the wild type, whereas *afp3* mutant plants exhibited a marked reduction in fruit set, indicating that *AFP3* positively influences this trait (**Fig. 6B**). However, despite the increased fruit set and higher number of fruits in *AFP3-OX* heterozygous plants, their overall yield of mature tomatoes was lower than that of wild-type plants (**Fig. 6C**). A similar trend was observed in *afp3* mutant plants, which also produced slightly reduced yields. While fruit ripening was comparable between wild-type and *afp3* mutant plants, it was severely delayed in *AFP3-OX* plants (**Fig. 6D**). Consequently, *AFP3-OX* plants produced more fruits that remained mostly smaller and unripe due to delayed maturation.

**Figure 6.**
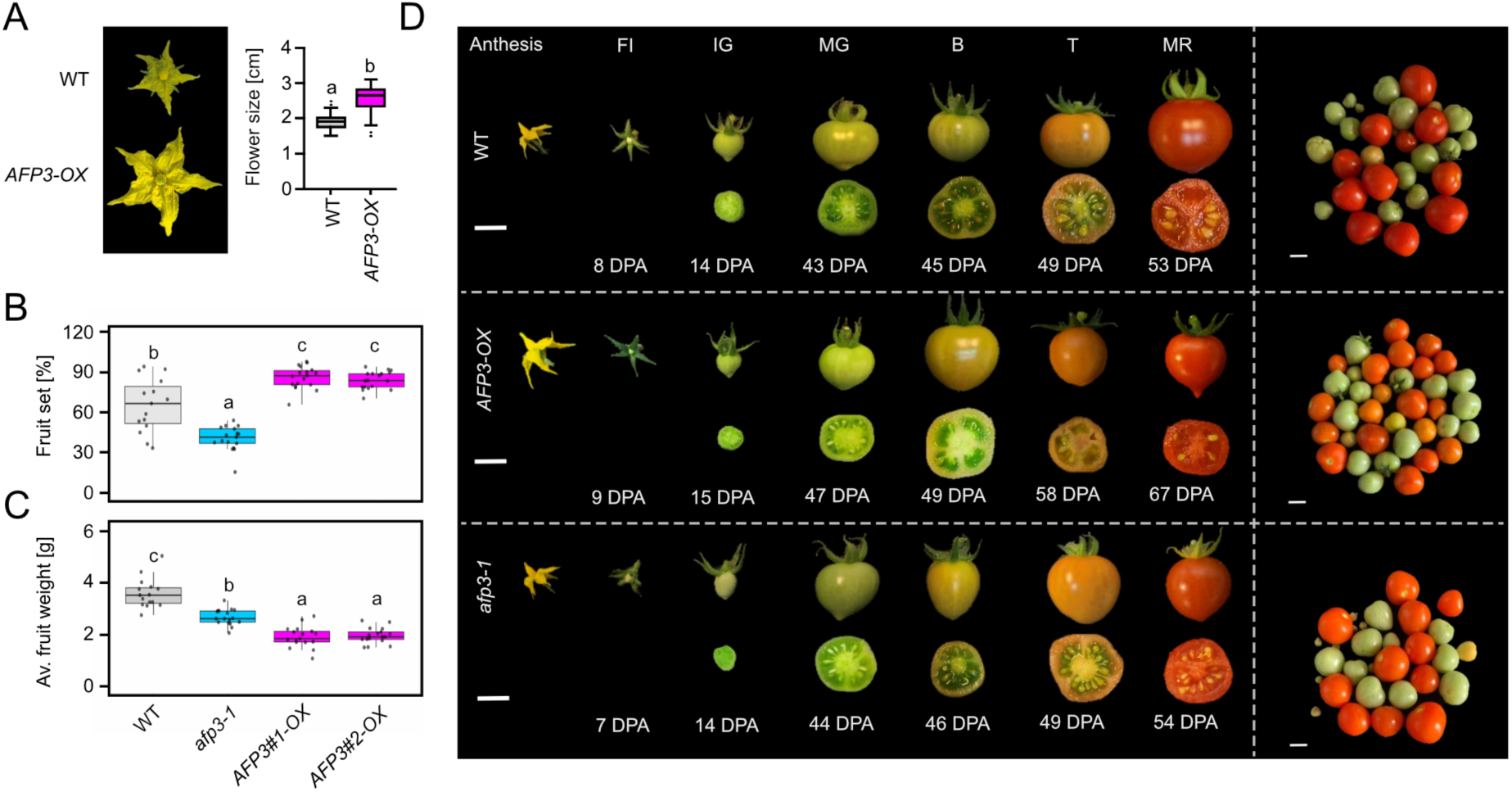
AFP3 and sAFP3 have synergistic and opposing function in flower and fruit development. **(A)** Picture showing the difference in size between an *AFP3-OX* flower and WT at anthesis stage (Scale bar= 1 cm). Included box plot showing the diameter length in cm of the flowers. One-way ANOVA with post hoc Tukey Test. Confidence level 0.95, significance alpha=0.05 (P < 0.05). Graphical representation of box plots in GraphPad Prism represents mean and SEM (flowers n=30 per genotype). **(B)** Total fruit set percentage per plant measured as number of fruits divided by the number of flowers in the plant. **(C)** Average weight of mature fruits at 130 Days After Germination (DAG), measured as weight of total mature tomatoes divided by the number. Scatter bar plots shows individual data points Mean and SEM (n= 15 control plants, n=15 to 21 mutant or transgenic plants). **(D)** Left panel: Pictures of fruit development from anthesis flowers to ripe stage (scale bar= 1cm) of the genotypes used for transcriptomic studies. Developmental stages (FI=Fruit Initiation; IG= Immature Green; MG: Mature Green; B= Breaker stage; T= Turning Orange Stage; MR= Mature Red stage) are measured as Days Post Anthesis (DPA). Right panel: Representative picture of tomatoes harvested at 130 DAG (scale bar= 1cm).

These findings suggest that both insufficient (*afp3* mutant) and excessive (*AFP3-OX*) expression of *AFP3* disrupt the delicate regulatory balance required for optimal reproductive development. In *afp3* mutants, the reduced expression likely impairs key processes in fruit development, directly limiting fruit set and yield. Conversely, overexpression of *AFP3* may enhance fruit initiation but leads to developmental delays and resource allocation imbalances, resulting in an increased number of immature or poorly developed fruits. This phenotypic trade-off highlights the importance of tightly regulated AFP3 activity, suggesting that precise spatial and temporal gene and protein activity is critical for maximizing fruit yield and quality. Thus, this comprehensive phenotypic analysis underscores the necessity of in-depth investigations into gene expression dynamics and their consequences across developmental stages and reproductive phases.

### Excessive AFP3 induces a physiological imbalance with reduced vigour and yield

As previously mentioned, establishing homozygote transgenic plants carrying the *AFP3-OX* transgene proved to be a challenge. We observed that some plants exhibited paler green coloration and stunted growth, and the resulting seeds from these plants often failed to germinate. As a result, we focused our efforts on healthy-looking heterozygote individuals. After several rounds of pooling seeds from homozygote individuals, we finally obtained enough seeds for further analysis. When compared to heterozygote individuals, we observed pale and dwarf plants that after genotyping turned out to be homozygotes (**Fig. 7A**). Gene expression analysis revealed that homozygote plants express *AFP3* at higher levels than heterozygotes (**Fig. 7B**). This finding suggests that AFP3 acts in a dose-dependent manner, where lower expression promotes growth and higher expression results in plants that are delayed in growth and appear less healthy.

**Figure 7.**
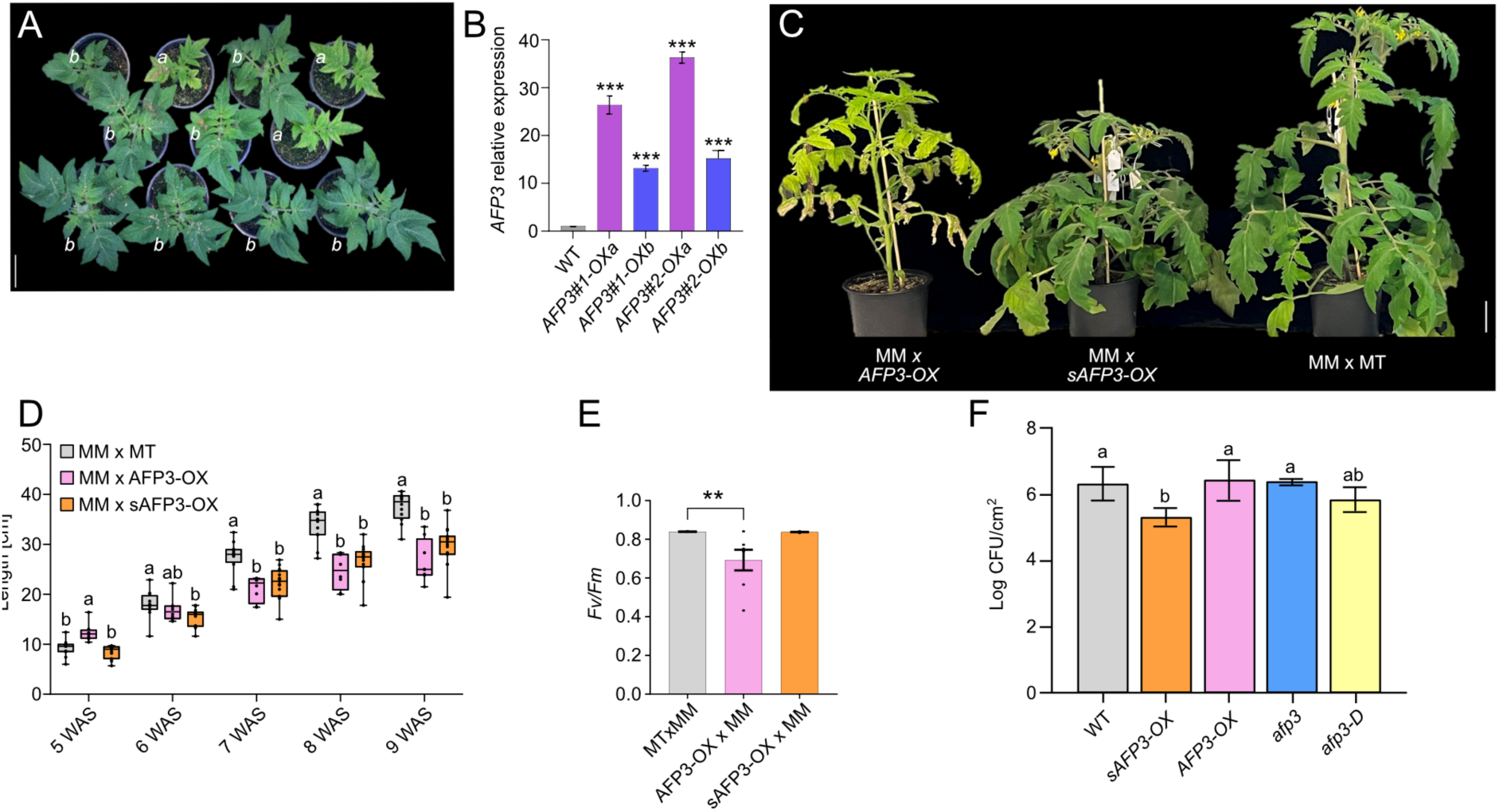
High levels of *AFP3* induce auto-immune responses. **(A)** Transgenic *AFP3-OX* plants that are homozygous *(a)* or heterozygous *(b)* 33 DAG. **(B)** RT-qPCR of the *AFP3* transcript from leaves of *AFP3-OX* homozygous (a) and heterozygous (b) plants. Panel shows higher relative expression of *AFP3* in homozygous compared to heterozygous plants. T-test (p-value * <0.05, ** <0.01, ***<0.001). **(C)** Crosses with the tomato Money Maker variety at 8 weeks after sowing (WAS) in soil. **(D)** Shoot length time course of the crossed lines from 5 to 9 WAS. One-Way ANOVA with post hoc Tukey test multiple comparison was performed for individual time points. **(E)** *Fv/Fm* ± SE (n=7-15) quantified by Handy PEA+ on uppermost leaves in 10 WAS Money Maker crosses. Student T-test (** P < 0.01). **(F)** Bacterial contents of plants infected with *Pseudomonas syringae*. One-Way ANOVA with post hoc Tukey test multiple comparison was performed for samples.

The tomato cultivar Micro-Tom harbors several loss-of-function mutations in genes involved in the brassinosteroid signaling pathway, resulting in its characteristic dwarf phenotype. Additionally, other mutations in Micro-Tom confer enhanced resistance to pathogens, making it a convenient and resilient model system for cultivation in space-limited environments. We were curious to what extent the genotype of the Micro-Tom variety influenced the phenotypes we observed. To address this question, we crossed the homozygote transgenic *AFP3-OX* and *sAFP3-OX* plants with the Moneymaker variety and as a control, we crossed Micro-Tom and Moneymaker wild-type plants. We expected the *sAFP3-OX* transgene to result in shorter offspring, while the *AFP3-OX* transgene was expected to lead to larger plants. Our analysis revealed that both crosses with the transgenic plants resulted in shorter plants. In the case of the *AFP3-OX* transgene, these plants were pale green and exhibited symptoms of stress and early senescence, while *sAFP3-OX* plants appeared darker green and healthy (**Fig. 7C**). A time-course analysis of plant height over a developmental period of nine weeks after sowing provided further evidence of the dwarf phenotypes observed. The smaller stature of the plants became apparent around seven weeks after sowing (**Fig. 7D**). Furthermore, pale green leaves and weak appearance observed in *AFP3-OX x MM* plants suggested defects in photosynthetic activity. To analyze the photosynthetic efficiency of these plants, we quantified the Maximum quantum efficiency of the photosystem II (PSII) photochemistry as Fv/Fm values. The analysis revealed that the PSII of the *AFP3-OX x MM* was strongly impaired compared with wild-type plants. While no differences were observed in *sAFP3-OXx MM* plants (**Fig. 7E**).

Given the stressed phenotype observed in *AFP3-OX* homozygotes and in heterozygous Moneymaker crosses, we investigated whether these visibly compromised plants might exhibit altered susceptibility to pathogen attack. To this end, we performed bacterial infection assays using *Pseudomonas syringae*. In these experiments, we consistently observed that transgenic plants overexpressing *sAFP3*, as well as *afp3-D* mutants, displayed slightly increased resistance to Pseudomonas infection (**Fig. 7F**). In contrast, *AFP3-OX* plants occasionally showed increased susceptibility in some assays; however, these results were inconsistent and the effects were modest. The similar phenotype observed in *sAFP3-OX* and *afp3-D* plants supports the conclusion that *afp3-D* is a dominant-negative allele that enhances pathogen resistance through a mechanism extending beyond native AFP3 function, and that this effect can be genetically engineered. These findings provide evidence that sAFP3 plays a role in suppressing growth and that AFP3 acts as a dose-sensitive regulator of development and senescence. Furthermore, the Micro-Tom cultivar displayed increased resistance to AFP3, tolerating higher levels of the gene dosage, while Moneymaker is more sensitive, exhibiting autoimmune defects even as heterozygote plants. These findings indicate that the genotype of the variety has a significant influence on the effect of AFP3.

## DISCUSSION

### Role of AFP3 and sAFP3 in plant development

In this study, we investigated the diverse functional roles of the tomato *AFP3* gene and its associated mRNA isoforms. The full-length *AFP3* isoform exhibited growth-promoting effects in the heterozygous state, whereas homozygous, high expression led to a debilitated phenotype. We also characterized its shorter isoform, which we named *sAFP3*, in regulating the growth and development of tomato plants. Recent advances in genomics have revealed that genes often express multiple alternative transcripts that may play critical roles under specific conditions or in response to different stimuli. Such transcripts often remain unannotated, and their biological significance elusive, especially in the context of plant responses to environmental stresses. Ectopic overexpression of the *AFP3* full-length isoform significantly reduced the number of axillary shoots of the corresponding transgenic plants, regardless of whether they were heterozygous or homozygous (**Suppl. Fig. S4**). These results are consistent with those obtained in Arabidopsis, where overexpression of *YFP-AFP2* also resulted in the formation of fewer lateral shoots^22^.

Transgenic plants homozygous for the *AFP3-OX* transgene and crosses of *AFP3-OX* to the Moneymaker variety resulted in stunted and diseased-looking plants that were paler green and underwent early senescence. Consistent with these phenotypes, we also observed reduced photosynthetic capacity in these plants (**Fig. 7E**). In *Nicotiana benthamiana* leaves, overexpression of *AFP2* has been shown to induce cell death through interactions with the SnRK1α1 subunit mediated by the B and C domains of the AFP2 protein^23,24^. SnRKs are Ser/Thr kinase complexes that are activated in the stress response to restore cellular energy homeostasis. Co-expression of ectopic SnRK1α1 antagonizes cell death, suggesting a role for this kinase in stress tolerance. We can speculate that the early signs of senescence and necrosis in *AFP3-OX* homozygous may result from an analogous mechanism where AFP3 inhibits kinases involved in stress regulation. In our RNA-seq data, we observe an overrepresentation of downregulated genes related to osmotic and salt stress (**Supplemental Dataset 1 and 2**). The finding that the A and B domains of AFP2 are necessary and sufficient for cell death induction suggests that sAFP3 alone cannot induce a stress phenotype because it only contains the C domain (**Fig. 1C**).

Like *sAFP3-OX*, the endogenous *sAFP3*, *afp3-D*, significantly inhibited plant growth. However, despite these strong phenotypes, transcriptomic analysis of *afp3-D* plants revealed minimal differences in gene expression levels. This is likely a result of *afp3-D* being expressed by its native *AFP3* promoter in the endogenous context, compared to the *sAFP3-OX* driven by a *35S* promoter. Consequently, *afp3-D* exerts its effects where and when it is naturally expressed, minimizing widespread transcriptional changes. Protein interaction studies, including yeast two-hybrid assays (**Fig. 4F**) and co-localization experiments, show that sAFP3 and AFP3 can interact in the nucleus (**Fig. 4B**). This interaction suggests a cooperative function in the genetic regulation of plant development. Given the high amino acid sequence conservation of sAFP3—particularly its similarity to the C-domain shared across the AFP protein family—there is significant potential to explore sAFP3 interactions with other AFP family members. However, unlike *sAFP3-OX* plants, where ectopic expression may influence multiple AFP proteins throughout development, the *afp3-D* variant is likely expressed predominantly in tissues where AFP3 is normally active. Importantly, because the endogenous *AFP3* gene has been disrupted by CRISPR-mediated deletion, the sAFP3 that is expressed in *afp3-D* can no longer interact with the native AFP3 protein. This targeted expression pattern likely restricts sAFP3 interactions to its intended molecular partners, minimizing unintended cross-talk with other AFP family members in unrelated developmental contexts. Such specificity may be advantageous for achieving precise modulation of developmental processes while avoiding off-target effects.

### Effect of AFP3 and sAFP3 on hormone-mediated signaling processes

Overexpression of *AFP3* promoted enhanced growth in the heterozygous state. In contrast, overexpression of *sAFP3* and its endogenously processed isoform (in *afp3-D*) resulted in decreased plant height (**Fig. 2A, B**). This reduction in growth is similar to that observed in the loss-of-function mutant *afp3*. We used the Micro-Tom cultivar for its compact size and fast life cycle, ideal for genetic studies. Its dwarf, bushy phenotype arises from mutations in the SELF-PRUNING (SP) and DWARF (D) genes, causing altered SP protein and truncated D protein, respectively^25,26^. Overexpression of *sAFP3* further enhanced the reduced growth phenotypes of the Micro-Tom tomatoes, which was transferable by crosses with the commercial Moneymaker variety (**Fig. 2 A, B** and **Fig. 7C, D**). AFP3, as a member of the AFP protein family, could potentially bind and inhibit ABI5 (ABA INSENSITIVE 5), the major transcriptional regulator of abscisic acid (ABA) signaling. Overexpression of AFP proteins generally results in a downregulation of ABA signaling through ABI5 inhibition^17^. This finding is also supported in tomato plants where sAFP3 (in *afp3-D*) caused an increase in ABA levels independently of AFP3 (**Fig. 5A**). The antagonistic relationship between abscisic acid (ABA) and brassinosteroids (BRs) in plant development is well established. BRs promote growth primarily through activation of the BRASSINAZOLE RESISTANT 1 (BZR1) transcription factor, which in turn downregulates *ABI5*, a key component of ABA signaling. In the Micro-Tom cultivar, which carries mutations in the BR signaling pathway, it is plausible that these alterations impact ABA signaling dynamics. Under such conditions, overexpression of an AFP protein that inhibits ABI5—such as in heterozygous *AFP3-OX* plants—could partially compensate for impaired BR signaling, thereby contributing to the observed increase in plant and flower size. Conversely, when *AFP3* is knocked out (*afp3*) or its function is competitively inhibited by sAFP3 overexpression (*sAFP3-OX*), the disruption of ABI5 repression may exacerbate the effects of compromised BR signaling. This could result in an exaggerated Micro-Tom phenotype, characterized by increased leaf distortion and a more rugose leaflet surface that we observed (**Suppl. Fig. S4D**).

The brassinosteroid (BR) deficiency characteristic of Micro-Tom plants is known to enhance gibberellin (GA) inactivation pathways, leading to reduced GA levels and smaller plant phenotypes^25^. In our transcriptomic analysis, we observed a predominance of downregulated genes, with Gene Ontology (GO) enrichment indicating suppression of biological processes related to growth, development, and gibberellin biosynthesis in both homozygous *AFP3-OX* and *sAFP3-OX* plants (**Supplementary Datasets 1 and 2**). In contrast, heterozygous *AFP3-OX* mutants displayed elongated shoots and a robust phenotype up to the late fruit development stage (**Fig. 2A, B**). These findings suggest that a high dosage of AFP3 may negatively influence GA biosynthetic processes. Interestingly, this interpretation is partially supported by our hormonal profiling, which revealed elevated GA levels in both *afp3* and *afp3-D* mutants (**Fig. 5B**).

### Transcriptional networks operating downstream AFP3 and sAFP3

Transcriptome analysis following the loss or ectopic expression of AFP3 and sAFP3 isoforms revealed widespread changes, including a substantial number of differentially expressed genes encoding transcription factors. This suggests that many of the observed developmental phenotypes may arise as secondary effects of altered expression of these regulatory genes. Some of these transcription factors may also be direct components of the AFP3-associated transcriptional complex, potentially participating in an autoregulatory feedback loop. However, the sheer number of transcription factors affected, particularly under *AFP3* overexpression, currently precludes comprehensive individual characterization. This complexity is especially evident in the *AFP3-OX* transcriptome, which includes more than 2,500 differentially expressed genes (DEGs) uniquely altered in these transgenic lines (**Fig. 3D**). These findings highlight the extensive pleiotropic effects of *AFP3* overexpression and suggest its involvement in modulating multiple signaling and developmental pathways.

In addition to the antagonistic functions of sAFP3 and AFP3, we also observed some degree of synergism. This synergism is reflected in the transcriptome dataset, as homozygous overexpression of either *AFP3* or *sAFP3* resulted in a large number of differentially expressed genes (DEGs) changing in a similar manner. In total, 284 genes were found to be downregulated when *AFP3* and *sAFP3* were overexpressed. Since several of these genes encode enzymes involved in amino acid biosynthesis, it seems plausible that the retarded growth of the transgenic plants could be a consequence of amino acid unavailability. In addition, hormones such as auxins are products of tryptophan metabolism and their deficiency could exacerbate the situation. Besides the down-regulated genes, we also found 197 genes that were up-regulated in the transgenic plants overexpressing either *AFP3* or *sAFP3*. However, no enriched gene ontologies were identified. It may be important to note that this gene set also includes the tomato *AFP1* gene, indicating a potential positive feedback loop. Furthermore, 22 of the 197 genes have an association with auxin, including 12 SAUR-like proteins. The latter may indicate increased auxin signaling in these transgenic plants despite the reduced growth of these plants.

### Role of AFP3 and sAFP3 in tissue differentiation

Loss of AFP3 or the presence of sAFP3 (as seen in *afp3-D* mutants or sAFP3-overexpressing plants) leads to reduced plant size, whereas heterozygous overexpression of AFP3 promotes growth (**Fig. 2**), highlighting their opposing roles in growth regulation. Molecularly, we identified connections between the AFP3/sAFP3 protein complex and growth regulation. Using affinity purification mass spectrometry (AP-MS), we identified a small gibberellic acid-stimulated Arabidopsis (GASA) protein as an interactor with both AFP3 and sAFP3. GASA proteins are small cysteine-rich proteins that have been shown to be transcriptionally upregulated in response to different hormone treatments and to be broadly involved in regulating organ growth ^27–29^. It thus appears plausible that sAFP3 limits the access of GASA proteins to the growth-promoting AFP3 protein complex, which hampers growth. This hypothesis is supported by the observation that GASA proteins can promote growth when AFP3 is limited.

The analysis of transgenic tomato plants with ectopic expression of either *sAFP3* or *AFP3* revealed both synergistic and antagonistic growth phenotypes. Using RNA-seq, we were also able to identify subsets of genes that were antagonistically expressed (**Fig. 3D**), although the numbers were relatively small. Interestingly, the subset of antagonistically regulated genes that is induced by AFP3 but repressed by sAFP3 contains a significant number of genes with functions in polarity regulation (**Table 1**). This is consistent with the observation that all AFP3 and sAFP3 transgenic plants, as well as the mutants generated in this study, exhibited smaller leaves and asymmetric leaf blades with leaflet torsions (**Suppl. Fig. S4D**). Among the differentially expressed genes is also a gene encoding gibberellic acid-20 oxidase (GA20OX). Furthermore, the leaf shapes observed in tomato plants closely resemble the overexpression of a GA20OX enzyme in tobacco^30^. These findings imply that the growth phenotypes we observed could be related to the de-regulation of gibberellic acid biosynthesis.

The finding that overexpression of AFP3 can cause dose-dependent phenotypic changes was surprising to us. A possible explanation is that ectopic overexpression of either AFP3 or sAFP3 always affects both endogenous AFP3 and sAFP3 transcripts. Thus, overexpression of AFP3 will lead to the formation of more AFP3 homodimers, while the endogenous sAFP3 will be titrated out. Conversely, overexpression of sAFP3 will trap most endogenous AFP3 proteins in AFP3/sAFP3 heterodimers, potentially inactivating AFP3. If all possible protein complexes (AFP3/AFP3, AFP3/sAFP3 and sAFP3/sAFP3) have independent as well as interdependent functions, overexpression of one isoform will always cause divergent effects by shifting the balance between the different complexes. This may explain why some effects on gene expression, but also on development, are synergistic.

### Effect of gene dosage and genotype on AFP3-induced stress responses

Ectopic expression of *AFP3* revealed a strong correlation between gene dosage on the phenotypic characteristics of transgenic plants. In the Micro-Tom cultivar, transgenic plants heterozygote for the *AFP3-OX* transgene showed growth promotion of vegetative tissues (**Fig. 2**). Homozygote *AFP3-OX* transgenic plants showed impaired growth (**Fig. 7** and **Suppl. Fig. S3**). These findings indicate a gene dosage effect, whereby the activity of the gene, the amount of mRNA it produces, and the abundance of protein correlate with the severity of the phenotypes observed. Gene dosage effects are well documented in the homeobox gene family, which in animal development often act as master regulators of the body plan, orchestrating the development along the anterior-posterior axis^31^. The *PAX6* gene, which plays an evolutionary conserved role in eye development, has been demonstrated to exhibit strong gene dosage effects. Reduced expression results in mild eye deformities, whereas excessive expression leads to severe, lethal craniofacial defects^32^. Our findings indicate that the *AFP3-OX* transgene causes severe growth retardation and early senescence when introduced into the Moneymaker background. This may suggest that AFP3 activity, likely lower in the Moneymaker background, becomes excessively elevated when combined with the *AFP3-OX* transgene, potentially surpassing a tolerable threshold of expression. Alternatively, one could speculate that the excessive expression of *AFP3* in homozygote Micro-Tom plants could lead to epigenetic modifications that are passed on to the Moneymaker cross. However, this is less probable, as we observed the growth-promoting effect of the heterozygote *AFP3-OX* transgene when we cross homozygote *AFP3-OX* plants with *afp3* mutant plants (not shown) in the Micro-Tom cultivar. Therefore, we conclude that gene dosage likely plays a critical role in the adverse phenotypes observed under high *AFP3* expression. Assuming that Moneymaker plants produce lower levels of the endogenous sAFP3 inhibitor, AFP3 overexpression may both suppress this inhibition and promote the accumulation of more active AFP3 protein. This suggests that the sAFP3 microProtein functions as an important regulatory element required to keep AFP3 activity in check, thereby maintaining proper developmental balance. Together, our results underscore the importance of tightly regulated *AFP3* expression for normal plant growth and development. These findings reveal a delicate interplay between gene dosage, cultivar background, and microProtein-mediated regulation. More broadly, this work highlights how quantitative shifts in repressor activity can have profound consequences on plant physiology, with potential implications for stress adaptation and developmental plasticity.

## MATERIALS AND METHODS

### Plant material and Growth conditions

Tomato plants *Solanum lycopersicum* cv. Micro-Tom (seed provider: BV Buzzy Thema) and Moneymaker (seeds provided by Marquardt S., Copenhagen) were utilized for phenotypic experiments. The plants designated for phenotypic analysis were cultivated under long day (LD) conditions, with temperatures maintained at 22°C and relative humidity set to 80% during the day, light intensity of 100 μmol m^-2^ s^-1^. At night, the conditions were adjusted to the same temperature but with a relative humidity of 65%. The seeds were initially sterilized using a solution of 70% ethanol and Tween 20 for 1 minute, followed by a 6% sodium hypochlorite solution diluted in water for 3 minutes. For germination seeds were kept in dark for 72 hours on ½ Murashige and Skooge (MS) 1% agar plates supplemented with Sucrose (30 g/L). Once the cotyledons emerged, the seedlings were transferred to soil mixed in a 1:3 ratio with vermiculite. Macronutrients and micronutrients were periodically added to the soil during watering.

### Generation of transgenic and CRISPR/Cas9 edited plants

Micro-Tom mutants were obtained using *Agrobacterium tumefaciens*-mediated transformation following the protocol by Fernandez *et al*.^33^. Overexpression lines (*OX-AFP3, OX-SAFP3*) were generated using oligos amplifying respectively the full-length CDS sequence and the second exon of *Solyc04g005380*. The products were cloned into the vector pJAN33 via gateway cloning. Plants were then transformed using the Agrobacterium strain GV3101 pMP90 RK and selected by BASTA (Glufosinate-Ammonium 9 mg/L) in tissue culture. Homozygous plants in T2 generation have been successively selected by BASTA treatment in soil and transcript levels were measured via RT-qPCR. Total RNA was extracted from leaves tissues from 4-5 weeks old plants, using Spectrum™ Plant Total RNA Kit (Sigma-Aldrich) and 1 µg purified RNA was used for reverse transcription using iScript ™ cDNA Synthesis Kit (BIO-RAD). RT-qPCR was performed using SYBR green (ThermoScientific) on a Biorad CFX384, using oligos against *AFP3* gene (FWD 5′-TGAGTGGAGGAGGAGGAGAA-3′ and REV 5-ACCACCTAATGACAACCCAAG-3′) for *AFP3-OX* lines and against the second exon of *AFP3* (FWD 5′-AGTGAGGCAAAAGCATTGGG-3′ and REV 5′- GCTTCACAAACTCAGCTGGT-3′) for *SAFP3-OX*lines. Gene expression was normalized using housekeeping gene GAPDH *Solyc05g014470* (FWD 5′-TGCTCCCATGTTTGTTGTGG-3′and REV 5′-CTCTTCCACCTCTCCAGTCC-3′). Relative expression was calculated with ΔΔCt method. Protein detection was carried out via Western blot on crude extracts material using Monoclonal ANTI-FLAG^®^ M2 antibody (Sigma-Aldrich). CRISPR/Cas9 mutants were generated using two sgRNAs (5′- GGAGATGGAGAATCTTTCATTGG-3′ and 5′-GGAAAGTGAGGCAAAAGCATTGG-3′) targeting the beginning of exon 1 and the beginning of exon 2. Both sgRNAs were cloned in the vector pKIR1.1 as described by Jin & Marquardt^34^ with a dual-sgRNAs cloning strategy. The middle border containing the sgRNAs scaffold was digested using BsaI. Transformation in plants was performed with use of Agrobacterium strain GV3101 pMP90 and selected by Hygromycin (35 mg/L). Regenerated plants surviving selection in plates were then genotyped using Edward method for the gDNA extraction^35^ and screened by PCR using the primers annealing in the 5′ and 3′UTRs (FWD 5′-AGATTTTTCCATTTGGGTGTGG-3′ and REV 5′- TGATGCATTTTGTGCCTCATG-3′). Plants showing a deletion were then sent for sequencing (Macrogene, Amsterdam) using PCR purified product via E.Z.N.A^®^ Cycle Pure Kit. Cas9-free plants were selected by PCR using primers annealing in the CAS9 gene (FWD 5′- GAAAGAAACTGGTGGACAGCAC-3′and REV 5′-CGTCGTAGGTGTCCTTGCTCAG-3′).

### Transgenic lines selection for phenotypic analysis

For the Micro-Tom phenotyping wild-type seeds were germinated in plates (described above) together with T3 *AFP3-1*, *afp3-D*, T2 *OX-AFP3-1/+, OX-AFP3-2/+,* T3 *OX-SAFP3-1, OX-SAFP3-2, OX-SAFP3-3* seeds, only plants germinated at the same time were selected for phenotyping. CRISPR/Cas9 homozygous and Cas9 free mutants were selected by PCR (described above). Overexpression homozygous lines were assessed by having 100% of survivors after BASTA treatment previous phenotyping analysis. *AFP3-OX* homozygous lines germination rate has decreased to 1-2% after time, showing in the few germinated plants a severe stress phenotype. Therefore, heterozygous plants were used for the phenotypic analysis, where 100 seeds were genotyped by PCR using primers specific to the BASTA gene (Forward: 5′-CAACCACGTCTTCAAAGCAA-3′, Reverse: 5′-AAGGATAGTGGGATTGTGCG-3′) and wild-type plants were excluded. 2-4 homozygous *AFP3-OX* plants germinated from the heterozygous pull showing the recurrent stress phenotype are shown independently in supplementary figures, because of too low number of plants for statistics.

For the Moneymaker phenotyping, crosses were conducted between the Micro-Tom wild type (WT) and two mutants (*OX-AFP3-2* and *OX-SAFP3-2*) with the Moneymaker variety serving as the female parent. Emasculation of Moneymaker flowers was performed to prevent self-fertilization, followed by pollination using pollen from Micro-Tom. After pollination, flowers were tagged and monitored until seed maturation. F1 seeds were harvested and then germinated in dark for 72h to promote uniform germination. Later on, seeds were sown in soil under LD at 22°C, 80% humidity (Materials and growth conditions described above). For genotypic verification of the F1 generation (*AFP3-OXx MM* and *SAFP3-OXx MM*), PCR was employed using primers specific to the BASTA gene (Forward: 5′-CAACCACGTCTTCAAAGCAA-3′, Reverse: 5′-AAGGATAGTGGGATTGTGCG-3′). To confirm the success of the crosses between the wild-type Micro-Tom and Moneymaker, these plants were cultivated alongside the original Micro-Tom and Moneymaker cultivars to evaluate phenotypic differences.

### 5′RACE deep-seq library preparation

For the stress treatments before the 5’ RACE procedure, seedlings were cultivated directly in liquid ½ Murashige and Skooge (MS) media enriched with Sucrose (30 g/L), in shaking conditions. Twelve Days After Imbibition (DAI), pulls of 10 seedlings were exposed to Flagelline at a concentration of 3.3 µM for 10 minutes to mimic biotic stress. Flg22 peptide with sequence Ac-QRLSTGSRINSAKDDAAGLQIA-OH (Schafer-N, www.schafer-n.com) with purity >95% was provided by Peter Brodersen, University of Copenhagen. Flg22 peptide was dissolved in dimethyl sulfoxide (DMSO) at 1 mg/mL (stock solution). Two more seedlings’ batches were treated respectively with 100 µM Abscisic Acid (ABA) for 4 hours, and NaCl at 120 mM for 4 hours to mimic abiotic stress conditions. Two control groups were included for comparison: one untreated and one exposed to DMSO alone. Plant material was collected in liquid Nitrogen and RNA extraction was performed using Spectrum™ Plant Total RNA Kit (Sigma-Aldrich). Using FirstChoice® RLM-RACE kit 10 µg of total RNA were processed performing a dephosphorylation with Calf Intestine Alkaline Phosphatase (CIP), and de-capping with Tobacco Acid Pyrophosphatase (TAP) treatment. CIP/Tap-treated RNA was then ligated to a 5′RACE Adapter and used for M-MLV Reverse Transcription. CIP/Tap-treated RNA was then ligated to a 5′RACE Adapter and used for M-MLV Reverse Transcription. 35 PCR cycles were run using the FWD 5′Outer primer 5’-GCTGATGGCGATGAATGAACACTG-3’ and a gene specific REV primer 5′-TGACAATGCTCAAATACCATTGCTCCATATTTC-3′. 1:10 of the PCR reaction was used as template for 35 cycles Nested PCR, using 5′Inner primer 5’- CGCGGATCCGAACACTGCGTTTGCTGGCTTTGATG-3’ and a gene specific REV primer 5′- GTTACTGTTTTGCATGATGCATTTTGTGCC-3′. The final PCR products were purified using E.Z.N.A^®^ Cycle Pure Kit and pulled together for Deep-sequencing (Novogene, UK). Paired-end raw data was submitted to Quality Check using FastQC to ensure data integrity before and after Trimming using Trimmomatic (Settings: leading=25, trailing=25, minlen=35, sliding window=4:25) (Bolger *et al*., 2014). High quality reads were converted in FASTA format. Using the “awk” command in the BASH Command Line in a Linux all the sequences containing the adaptor sequence 5′-GAACACTGCGTTTGCTGGCTTTGATG-3′ (provided by the RACE kit) were kept. The extracted reads were mapped to the reference WT gene sequence, using the minimap2 tool on the Galaxy platform. BAM files were analyzed and visualized in IGV (Integrative Genomics Viewer).

### Tissue specific qRT-qPCR

Samples for tissue specific analysis were harvested using Liquid Nitrogen and RNA extracted using RNeasy Plant Mini Kit (Qiagen). 1 µg purified RNA was used for reverse transcription using iScript ™ cDNA Synthesis Kit (BIO-RAD). RT-qPCR was performed using SYBR green (ThermoScientific) on a Biorad CFX384, using oligos against AFP3 (exon 1) (FWD 5′-TGAGTGGAGGAGGAGGAGAA-3′and REV 5′-ACCACCTAATGACAACCCAAG-3′) and against the exon 2 of AFP3 (FWD 5′-GACATGCCATGTGTTTTCGC -3′and REV 5′- GCTTCACAAACTCAGCTGGT-3′). The absolute expression with ΔΔCt method was calculated and normalized with the housekeeping gene ACTIN *Solyc11g005330* (FWD 5′- GGTCGTACAACTGGTATTGTG-3′and REV 5′-TAAATCACGACCAGCAAGATCC-3′). 3-4 biological replicates were used for each sample, for small amounts of tissues like seedlings, 6 DAG roots, stems and leaves, and flowers, 3 plants were used for biological replicate. Plants were grown in LD conditions.

### Subcellular localization

Constructs for subcellular localization in 5-week-old *Nicotiana benthamiana* leaves were prepared by cloning the *AFP3* and *SAFP3* CDS sequences into the Gateway vector pK7AFP3F2. This vector includes an eGFP tag at the N-terminus, under the control of the CaMV 35S promoter. An empty vector was utilized as a control. These constructs were co-infiltrated along with a vector carrying the p19 silencing suppressor; p19 alone was infiltrated as a negative control. Transient expression was achieved using *Agrobacterium tumefaciens* GV3101 cultures at an optical density (OD) of 0.3, which were incubated in Infiltration Buffer (10 mM MgCl2, 10 mM MES, pH 5.6, adjusted to pH 7.2) with 0.2 mM acetosyringone for 4 hours. The analysis was conducted on three plants infiltrated per construct, three days post-infiltration, employing the Leica Stellaris 8 confocal microscope. Leaf disks were excited with a 470 nm pulsed laser (10 MHz), and emissions from 500 nm to 560 nm were recorded. Images were captured using a 40x objective and analyzed with the LasX tool.

### Co-localization

For subcellular localization, the coding sequences of *AFP3* and *SAFP3* were cloned into the Gateway vectors *pK7AFP3F2* and *pEarlyGate104*, the latter of which contains the mCherry tag at the N-terminus and is under the control of the *CaMV35S* promoter (provided by Sabine Müller/Dorothee Stöckle, ZMBP Tübingen). Transient expression in *Nicotiana benthamiana* leaves was performed as described above, using three plants as individual biological replicates. Epidermal cells were imaged after three days using a Leica Stellaris 8 confocal microscope. Samples were excited with a 470 nm pulsed laser (10 MHz), and emissions from 500 to 560 nm were recorded. Images were analyzed using the LasX tool.

### Transcriptomic library preparation

For the RNA-seq library preparation, Micro-Tom 6 DAG seedlings were grown in ½ Murashige and Skooge (MS) 1% agar plates supplemented with Sucrose (30 g/L). For each genotype 3 biological replicate (3 seedlings per replicate were used). The libraries include wild-type, homozygous *OX-AFP3-2*, homozygous *OX-SAFP3-2*, homozygous *afp3-D*and homozygous *AFP3-1*. Plant material was collected and grinded in liquid Nitrogen, RNA extraction performed with Spectrum™ Plant Total RNA Kit (Sigma-Aldrich). RNA quality and concentration were assessed using a NanoDrop spectrophotometer and Qubit. RNA was sequenced on an Illumina HiSeq platform by Novogene (15G), and bioinformatics analysis was carried out by UPSC Bioinformatics Facility.

### RNA-Seq preprocessing

Raw sequencing reads were filtered for residual ribosomal RNA (rRNA) contamination by using SortMeRNA^36^ (v4.3.4; --fastx--sam --num_alignments 1) and the rRNA sequences provided with SortMeRNA (rfam-5s-database-id98.fasta, rfam-5.8s-database-id98.fasta, silva-arc-16s-database-id95.fasta, silva-bac-16s-database-id85.fasta, silva-euk-18s-database-id95.fasta, silva-arc-23s-database-id98.fasta, silva-bac-23s-database-id98.fasta and silva-euk-28s-database-id98.fasta). Non-rRNA reads were trimmed for sequencing adaptors and filtered for quality by using Trimmomatic^37^ (v0.39; settings TruSeq3-PE-2.fa:2:30:10 LEADING:3 SLIDINGWINDOW:5:20 MINLEN:50). Read quality was assessed before and after rRNA removal and quality filtered by FastQC (http://www.bioinformatics.babraham.ac.uk/projects/fastqc/). Filtered reads were pseudo-aligned to Solanum lycopersicum transcriptome (ITAG4.0, obtained from Phytozome v13) using Salmon^38^ (v1.9.0; non default settings: --gcBias --seqBias). In separated analyses where the actual alignment was needed, STAR^39^ (v2.7.10a; settings --alignIntronMax 11000) was used to align filtered reads onto Solanum lycopersicum genome (ITAG4.0). The filtered reads generated were visualized on IGV_2.17.3.

### DEG analysis

Per-gene read counts from Salmon were imported in R (v4.3.1) and normalized using a variance stabilizing transformation as implemented in DESeq2 ^40^ (v1.42.1). Similarly, within the biological replicates (e.g. Principal Component Analysis (PCA) was assessed by using custom R scripts, available from https://github.com/nicolasDelhomme/tomato-AFP3-LNJ. Differential expression was performed using DESeq2 with FDR adjusted p-values threshold at 0.01. The raw data has been submitted to Gene Expression Omnibus (GEO) and is available under accession number XXX. Heatmap showing the DEGs in all genotypes were prepared by pheatmap package in R. To focus on the biological significance of the DEGs, Gene Ontology Resource (https://geneontology.org) was used using *Solanum lycopersicum* genome reference.

### AP-MS and proteomic analysis

For the Affinity Purification-Mass Spectrometry study (AP-MS), seedlings of *OX-AFP3-2* and *OX-SAFP3-2*, as well as wild-type and T0 *OX-eGFP* plantlets were grown in ½ Murashige and Skooge (MS) 1% agar plates supplemented with Sucrose (30 g/L) in LD. A pull of 10-15 seedlings were collected in liquid nitrogen and grinded with mortar and pestle. For protein extraction and purification grinded samples (> 2g) were resuspended in an equivalent volume of SII buffer (comprising 100 mM NaPhosphate pH 8.0, 150mM NaCl, 5mM EDTA, 5mM EGTA, 0.1% TX-100) supplemented with EDTA-free protease inhibitor cocktail, 1mM PMSF(phenil metasulphonate), 1x phoSTOP and MG132 5uM) and incubated with rotation for 10 min at 4°C. The samples were sonicated for 10 seconds each (0.5 cycles on/off) at 10% power followed by three cold clarification steps for clarification. For the immunoprecipitation of 3XFLAG-tagged proteins, the clarified lysates were incubated with cross-linked anti-FLAG M2 beads (Sigma), at a ratio of 10µl beads/1mg of protein extract, for 1h at 4°C, accordingly to the manufactureŕs guidelines. The beads were then washed thrice with SII buffer and subsequently three times with a freshly prepared 25mM Ammonium Bicarbonate solution. Following these wash steps, the beads were rinsed twice with PBS and immediately frozen in liquid Nitrogen. Protein-Protein Interaction analysis via mass spectrometry and bioinformatics were conducted by Novo Nordisk Foundation Center for Protein Research’s Proteomics Research Infrastructure, utilizing 3XFLAG-SAFP3 and 3XFLAG-AFP3 as bait proteins (Baits1 and Baits2), and 3XFLAG-eGFP and wild-type samples as controls (control1 and control2). Data analysis, including the statistical assessment of protein expression levels, was executed using a custom Python script based on methodologies developed from the Clinical Knowledge Graph’s automated analysis pipeline ^41^. Proteins deemed potential contaminants, indicated by matches to a reverse decoy database or identified solely by modified sites, were excluded. Protein intensities were transformed to logarithmic scale (log2) and only those proteins with at least 2 valid measures in at least one group were filtered out. Exclusive proteins for each experimental group were identified requiring complete absence in control samples and presence in at least 60% of the experimental samples. Missing values were imputed using the MinProb method (width=0.2 and shift=1.8 ^42^). Differentially expressed proteins were identified via unpaired t-tests with p-values adjusted (for multiple hypothesis) using both Benjamini-Hochberg and permutation-based FDR corrections ^43^. Significant proteins, defined by an FDR of 0.01, were visualized in Volcano plots created with GraphPad Prism, highlighting those with a log2 fold-change greater than 2 and a p-value less than 0.01. For the identification of specific interactors of the Baits (*SAFP3-OX*and *OX-AFP3*) proteins detected also in *OX-eGFP* and WT were excluded to ensure specificity. A comprehensive list of common interactors for both *AFP3-OX* and *SAFP3-OX* was compiled using R studio, systematically excluding proteins uniquely interacting with each. Venn diagram was made using Ghent University Tool (https://bioinformatics.psb.ugent.be/webtools/Venn/).

### Yeast Two-Hybrid

Constructs for the Y2H assay were prepared via gateway cloning as AFP3 and SAFP3 CDS were combined into pGBKT7-GW and pGADT7-GW vectors. The constructs were transformed into AH109 yeast strain via PEG-LiAc mediated sequential transformation (Yeast Protocols Handbook, Clonetech). During primary transformation, the positive yeast colonies on SD single dropout medium plates lacking either leucine(L) or tryptophan(W) were chosen and proceeded for secondary transformation. The positive colonies from secondary transformation were allowed to grow on SD/-LW media. After 3-5 days the positive colonies on SD/-LW were selected and grown on both SD/-LW and SD/-LWH media by drop assay to visualize interaction. For each set of AFP3 and SAFP3 constructs, the protein-protein interactions were further confirmed in their reciprocal combinations. The experiment used empty pGBKT7-GW x pGADT7-GW as negative and pGADT7-TOPLESS x pGBKT7-miP1a as positive controls.

### Hormone Profiling (Hormonomics)

Hormone profiling was performed using 30-day-old leaf tissue collected from three tomato genotypes: WT (Micro-Tom), *afp3-1*, and *afp3-D*. The analysis was conducted by the Swedish Metabolomics Centre (www.swedishmetabolomicscentre.se), Umeå, Sweden. Approximately 20 mg of frozen leaf material per sample was homogenized using three carbide beads in a bead mill (27 Hz, 10 min, 4 °C; MixerMill, Retsch GmbH, Haan, Germany) and extracted in 10 mL of ice-cold 60% aqueous acetonitrile (v/v) containing a mixture of ^13C-or deuterium-labeled internal standards. The homogenates were centrifuged at 14,000 rpm for 10 min at 4 °C. Supernatants were purified using solid-phase extraction (SPE) on Oasis HLB columns (30 mg, 1 cc; Waters Inc., Milford, MA, USA) pre-conditioned with 1 mL of 100% methanol followed by 1 mL of Milli-Q water. The flow-through was collected during elution with 0.5 mL of 30% aqueous acetonitrile. Samples were evaporated to dryness using a SpeedVac concentrator (SpeedVac SPD111V, Thermo Scientific, Waltham, MA, USA) and reconstituted in 10 µL acetonitrile followed by 40 µL water. Extracts were transferred to insert-equipped vials for LC-MS analysis. Analysis was carried out on an Agilent 1290 Infinity Binary UHPLC system coupled to a 6495D Triple Quadrupole MS/MS System with Jet Stream and Dual Ion Funnel technologies (Agilent Technologies, Santa Clara, CA, USA). Chromatographic separation was achieved using an Acquity UPLC CSH C18 column (150 mm × 2.1 mm, 1.7 μm; Waters) maintained at 40 °C. A 10 μL injection volume was used. The mobile phases were 0.01% formic acid in water (A) and 0.01% formic acid in acetonitrile (B) with a flow rate of 0.25 mL min⁻¹. The gradient was as follows: 5% B for 10 min, ramped to 80% B over 20 min, held at 80% for 1 min, decreased back to 5% in 0.5 min, and equilibrated for 2.5 min. Multiple reaction monitoring (MRM) transitions were optimized using MassHunter MS Optimizer (Agilent Technologies) and compound-specific parameters are listed in Supplementary Table X. Quantification was performed using Agilent MassHunter Quantitative Analysis software (version 12.0). Hormone concentrations were determined based on the ratio of each endogenous compound to its corresponding stable isotope-labeled internal standard and normalized to sample fresh weight (pmol/mg). This analysis targeted multiple phytohormones, including auxins, cytokinins, gibberellins, abscisic acid, jasmonates, salicylic acid, and brassinosteroids, as described in Šimura et al. (2018) ^44^, with minor modifications. Peaks were integrated and processed using Agilent MassHunter Quantitative Analysis (for QQQ, version 12.0). Concentrations were calculated from the ratio of endogenous metabolite and deuterated or 13C-labeled standards, compared to a calibration curve of increasing endogenous standard. Concentrations were calculated as pmol per mg sample.

### Dark-adapted chlorophyll fluorescence measurements

Chlorophyll fluorescence was measured on uppermost fully expanded leaves of Moneymaker crosses (n plants= 7 to 15 per genotype) using the Hansatech Handy-PEA+ fluorimeter (Hansatech, UK). Leaves were dark adapted for 30 minutes and exposed to light pulse for 1 s, intensity 3500 µM, gain x01. The maximum quantum yield of photosystem II (PSII) was measured and *Fv/Fm* values plotted as scatter plot bar in GraphPad Prism.

## ACKNOWLEDGEMENTS

We acknowledge the funding from NovoCrops Centre (Novo Nordisk Foundation project number 2019OC53580 to S.W.), the Independent Research Fund Denmark (0136-00015B and 0135-00014B to S.W.), the Novo Nordisk Foundation (NNF18OC0034226 and NNF20OC0061440 to S.W.), Vetenskapsrådet (Reg. No. 2024-04792 to S.W.) and the European Research Council (ERCSynG RESYDE No. 101167406 to S.W.). We thank the University of Copenhagen Protein Research Infrastructure (PRI) for proteomics experiments and Dr. Tran Thien Long for critically reading the manuscript.

---

**Supplementary Figure S1.**
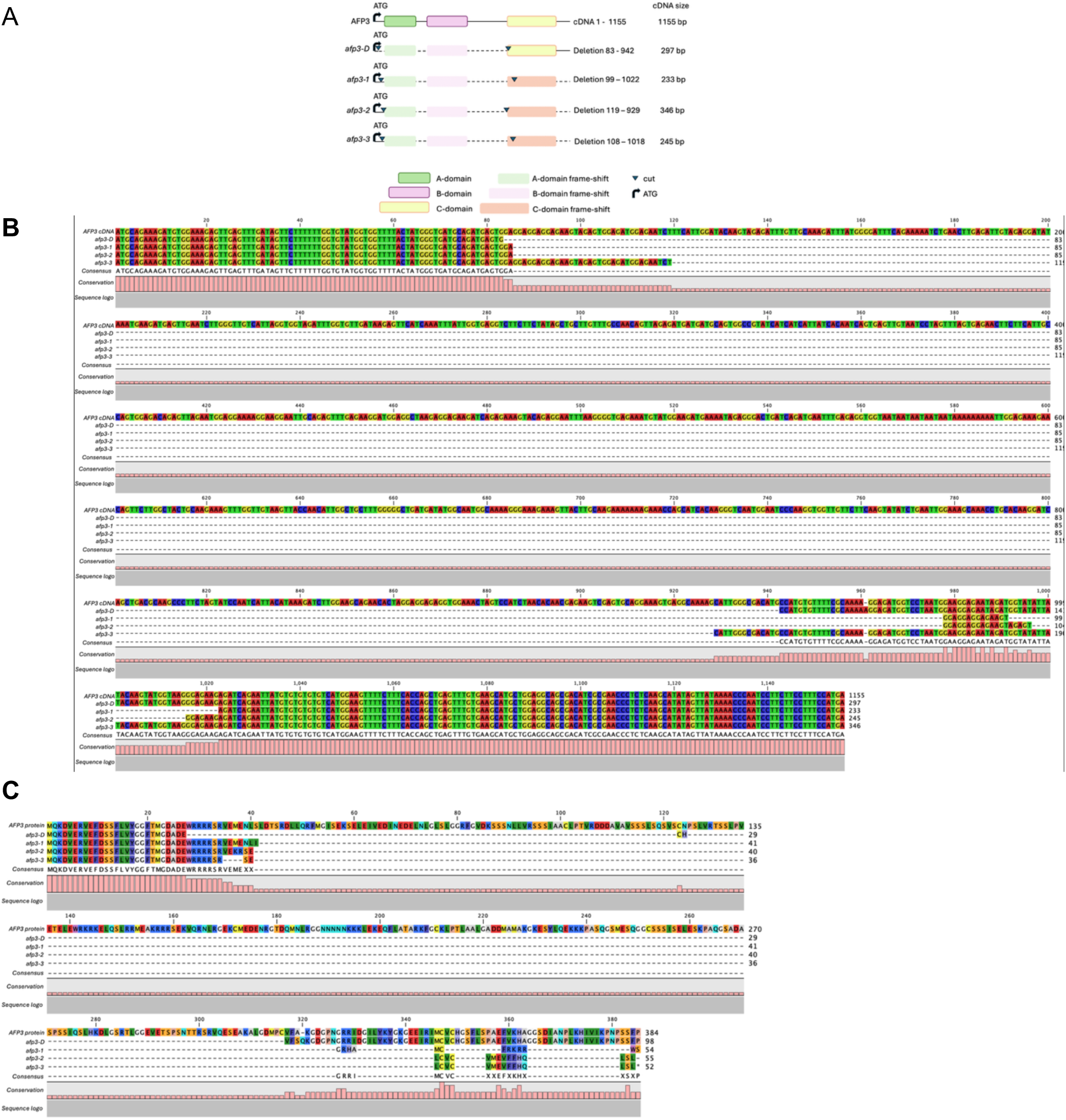
Characterization of *afp3* CRISPR lines. (**A**) Schematic representation of the *afp3-D* (gain-of-function), *afp3-1*, *afp3-2*, and *afp3-3* (loss-of-functions) CRISPR lines. The precise CRISPR-induced nucleotide deletions are indicated with reference to the *AFP3* cDNA sequence, along with the resulting cDNA length after deletion. (**B**) Alignment of Sanger-sequenced cDNAs from the CRISPR mutants. (**C**) Predicted amino acid sequences derived from the mutant cDNAs shown in (B), highlighting the in-frame /frame-shift deletions and their effect on the resulting protein products.

**Supplementary Figure S2.**
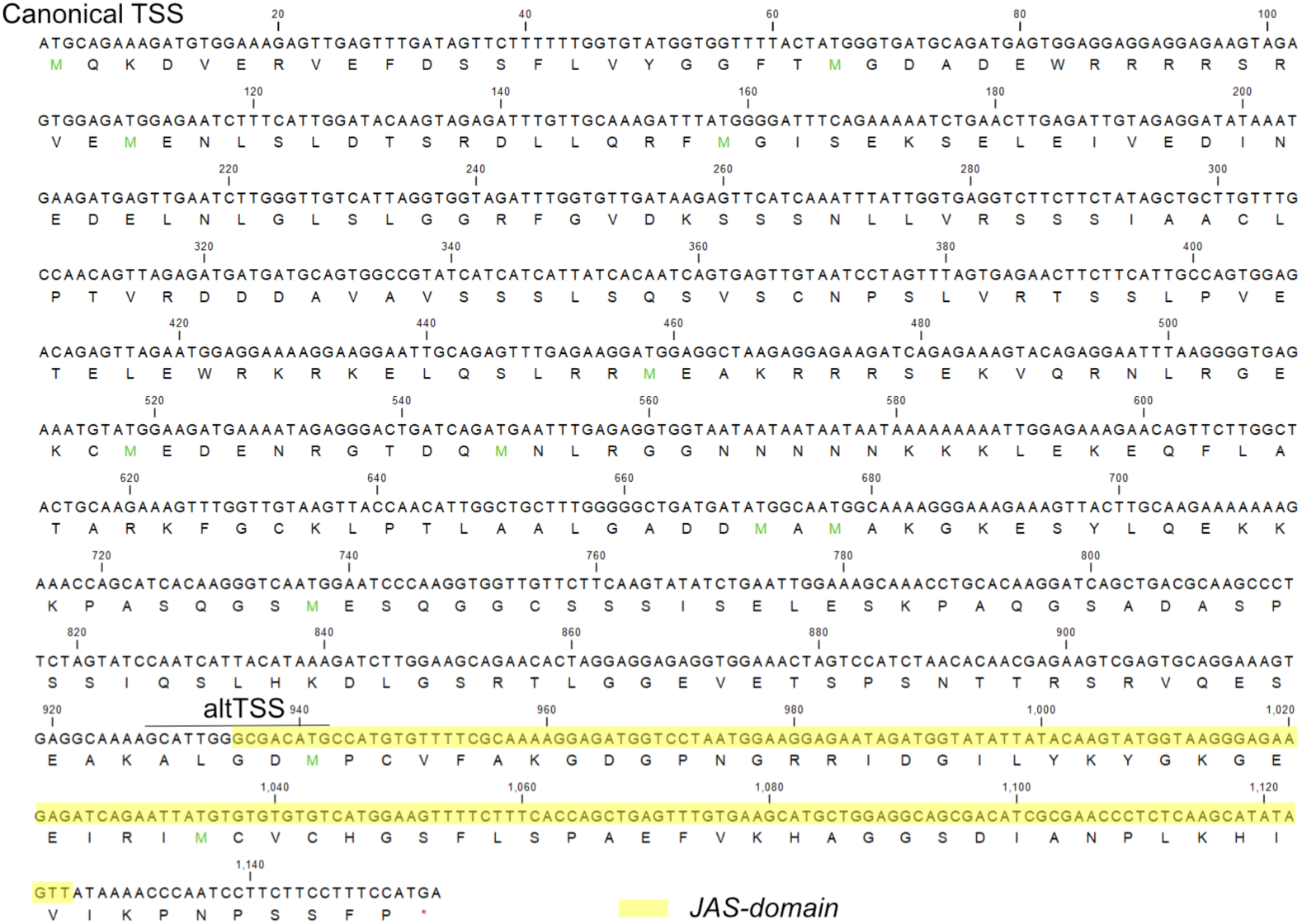
CDS sequence of *AFP3* gene and below translated +1 ORF. In the figure is shown the canonical TSS, represented by the already annotated ATG on NCBI and SolGenomics databases. We show the alternative TSS detected by 5′RACE Deep-seq experiment. In yellow, we highlight the predicted JAS domain according to the Protein domain prediction tool.

**Supplementary Figure S3.**
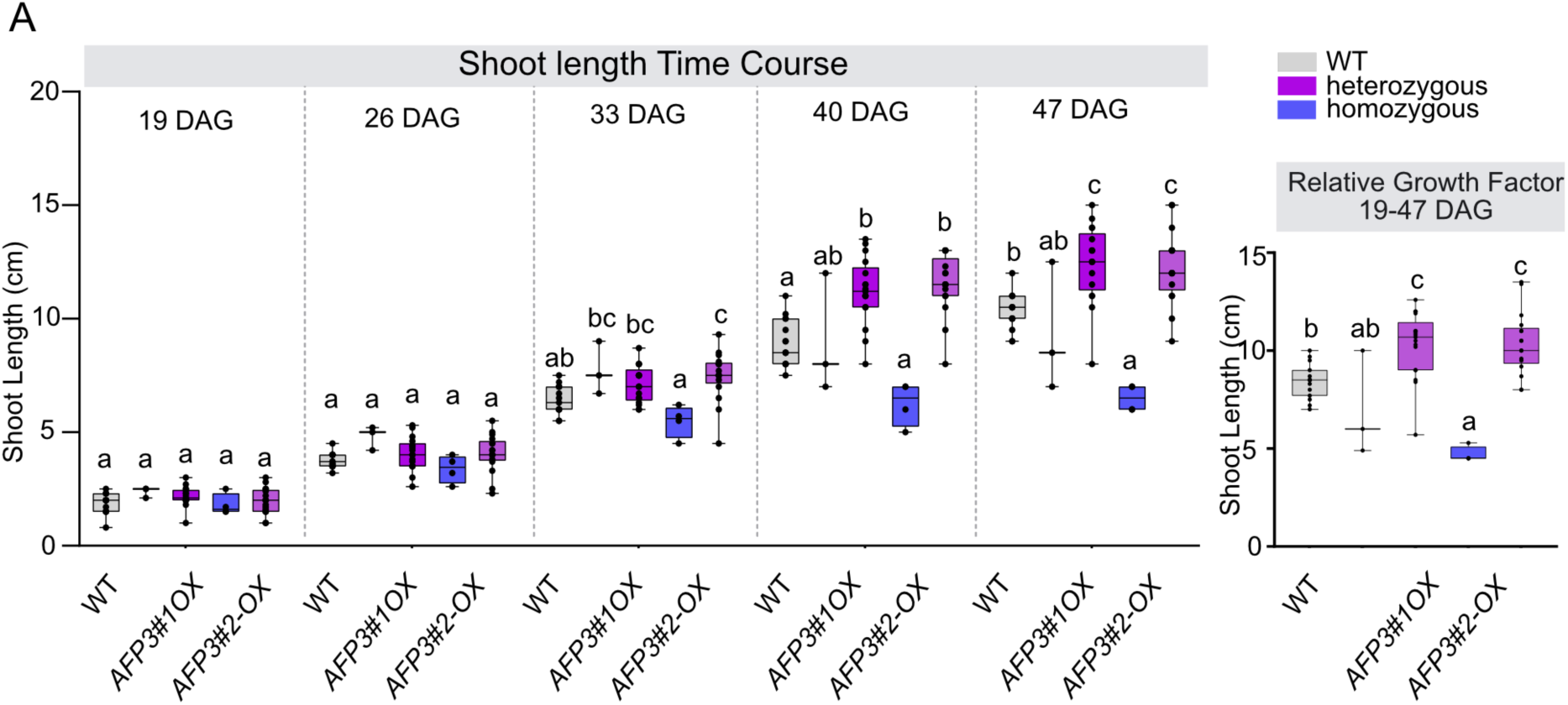
(A) Shoot Length measures represented as a Time course of 19, 26, 33, 40 and 47 DAG. The analysis compares WT (grey), homozygous *AFP3#1-OX* and *AFP3#2-OX* (blue), and heterozygous *AFP3#1-OX /+ and AFP3#2-OX /+* (purple); **(B)** Graphic representation of Plantś shoot growth as Relative Growth between 19 and 47 Days After Germination. One-way ANOVA with post hoc Tukey Test multiple comparisons for individual time points is made in R studio. Box plot is made in GraphPad Prism represents individual data points in addition to mean + SEM (n= 15 control plants, n=15 heterozygous plants, n=2-4 homozygous plants).

**Supplementary Figure S4.**
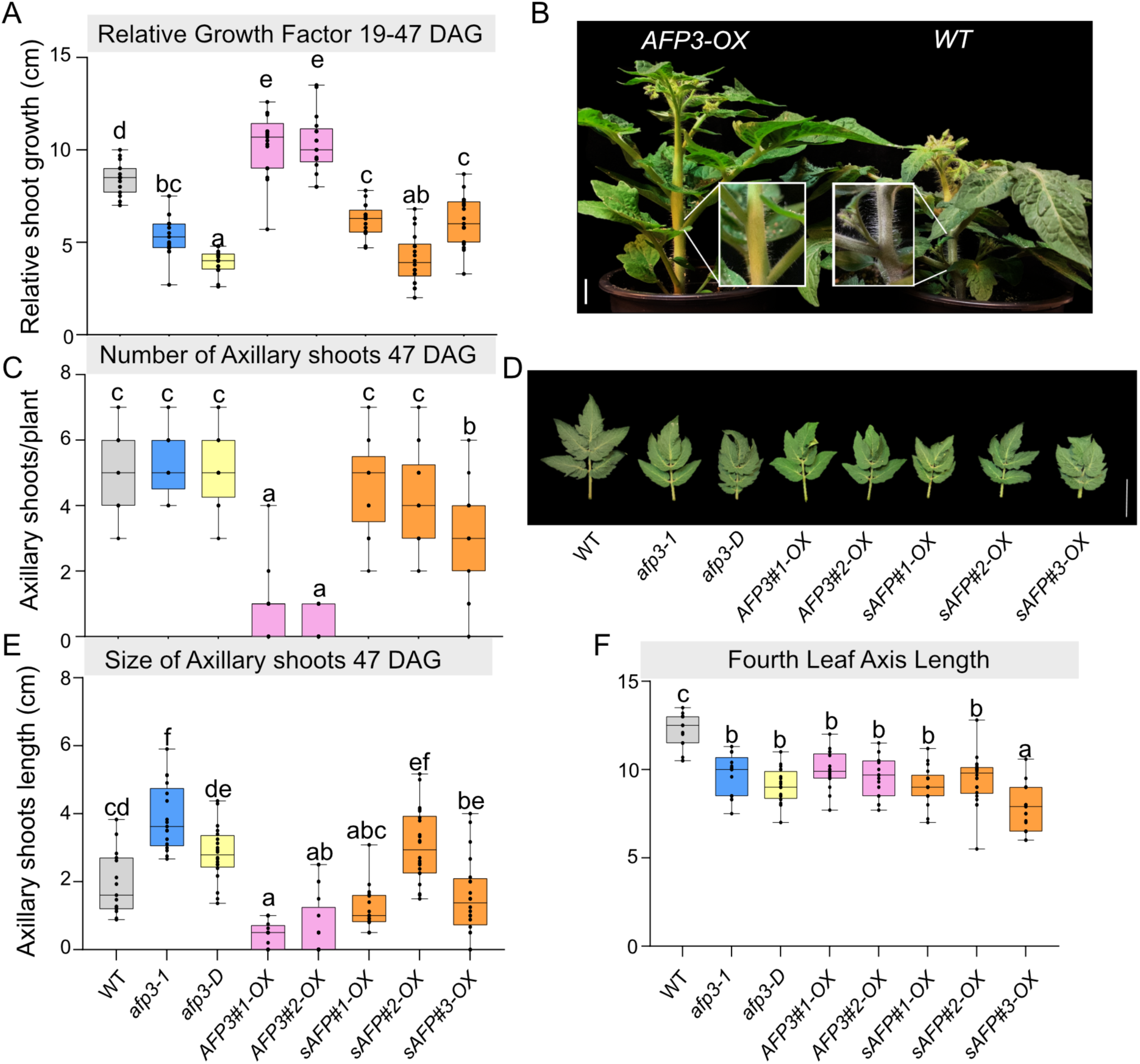
(A) Graphic representation of plants shoot growth as Relative Growth between 19 and 47 Days After Germination. **(B)** Number of Axillary shoots emerged on the main stem at 47 DAG per plant. **(C)** Size in cm of Axillary shoots measured as mean per plant at 47 DAG. Statistics made with One-Way ANOVA with post hoc Tukey Test in R studio. Confidence level 0.95, significance alpha=0.05 (P < 0.05). Graphical representation of box plots in GraphPad Prism represents individual data points in addition to mean + SEM (n= 15 control plants, n=15 to 21 mutant plants). **(D)** Image of plants at 47 DAG showing with a magnification of the main photo, no axillary shoot growth in *AFP3-OX* plants compared to WT. Scale bar = 1 cm. **(E)** Image showing Fourth leaf starting from bottom (not including cotyledons) at 33 DAG of the lines included in the phenotype. Image shows leaves of the different lines to be smaller and more curled compared to the WT (bar scale=5 cm). **(F)** Plot of the main axis length of the Fourth leaf at 33 DAG. Statistics made with One-Way ANOVA with post hoc Tukey Test in R studio. Confidence level 0.95, significance alpha=0.05 (P < 0.05). Graphical representation of box plots in GraphPad Prism represents individual data points in addition to mean + SEM (n= 15 control plants, n=15 to 21 mutant plants).

## REFERENCES

1. Reddy, A.S., Marquez, Y., Kalyna, M., and Barta, A. (2013). Complexity of the alternative splicing landscape in plants. Plant Cell 25, 3657–3683. 10.1105/tpc.113.117523.

2. Kashkan, I., Timofeyenko, K., and Růžička, K. (2022). How alternative splicing changes the properties of plant proteins. Quant Plant Biol 3, e14. 10.1017/qpb.2022.9.

3. Tognacca, R.S., Rodríguez, F.S., Aballay, F.E., Cartagena, C.M., Servi, L., and Petrillo, E. (2023). Alternative splicing in plants: current knowledge and future directions for assessing the biological relevance of splice variants. J Exp Bot 74, 2251–2272. 10.1093/jxb/erac431.

4. Ruiz-Orera, J., Villanueva-Cañas, J.L., and Albà, M.M. (2020). Evolution of new proteins from translated sORFs in long non-coding RNAs. Exp Cell Res 391, 111940. 10.1016/j.yexcr.2020.111940.

5. Lauressergues, D., Couzigou, J.M., Clemente, H.S., Martinez, Y., Dunand, C., Bécard, G., and Combier, J.P. (2015). Primary transcripts of microRNAs encode regulatory peptides. Nature 520, 90–93. 10.1038/nature14346.

6. Chen, Y., Ho, L., and Tergaonkar, V. (2021). sORF-Encoded MicroPeptides: New players in inflammation, metabolism, and precision medicine. Cancer Lett 500, 263–270. 10.1016/j.canlet.2020.10.038.

7. Merino-Valverde, I., Greco, E., and Abad, M. (2020). The microproteome of cancer: From invisibility to relevance. Exp Cell Res 392, 111997. 10.1016/j.yexcr.2020.111997.

8. Thieffry, A., López-Márquez, D., Bornholdt, J., Malekroudi, M.G., Bressendorff, S., Barghetti, A., Sandelin, A., and Brodersen, P. (2022). PAMP-triggered genetic reprogramming involves widespread alternative transcription initiation and an immediate transcription factor wave. Plant Cell 34, 2615–2637. 10.1093/plcell/koac108.

9. Hong, S.Y., Sun, B., Straub, D., Blaakmeer, A., Mineri, L., Koch, J., Brinch-Pedersen, H., Holme, I.B., Burow, M., Lyngs Jørgensen, H.J., et al. (2020). Heterologous microProtein expression identifies LITTLE NINJA, a dominant regulator of jasmonic acid signaling. Proc Natl Acad Sci U S A 117, 26197–26205. 10.1073/pnas.2005198117.

10. Lopez-Molina, L., Mongrand, S., Kinoshita, N., and Chua, N.H. (2003). AFP is a novel negative regulator of ABA signaling that promotes ABI5 protein degradation. Genes Dev 17, 410–418. 10.1101/gad.1055803.

11. Garcia, M.E., Lynch, T., Peeters, J., Snowden, C., and Finkelstein, R. (2008). A small plant-specific protein family of ABI five binding proteins (AFPs) regulates stress response in germinating Arabidopsis seeds and seedlings. Plant Mol Biol 67, 643–658. 10.1007/s11103-008-9344-2.

12. Finkelstein, R.R., and Lynch, T.J. (2000). The Arabidopsis abscisic acid response gene ABI5 encodes a basic leucine zipper transcription factor. Plant Cell 12, 599–609. 10.1105/tpc.12.4.599.

13. Chang, G., Yang, W., Zhang, Q., Huang, J., Yang, Y., and Hu, X. (2019). ABI5-BINDING PROTEIN2 Coordinates CONSTANS to Delay Flowering by Recruiting the Transcriptional Corepressor TPR2. Plant Physiol 179, 477–490. 10.1104/pp.18.00865.

14. Krogan, N.T., Hogan, K., and Long, J.A. (2012). APETALA2 negatively regulates multiple floral organ identity genes in Arabidopsis by recruiting the co-repressor TOPLESS and the histone deacetylase HDA19. Development 139, 4180–4190. 10.1242/dev.085407.

15. Oh, E., Zhu, J.Y., Ryu, H., Hwang, I., and Wang, Z.Y. (2014). TOPLESS mediates brassinosteroid-induced transcriptional repression through interaction with BZR1. Nat Commun 5, 4140. 10.1038/ncomms5140.

16. Lynch, T., Née, G., Chu, A., Krüger, T., Finkemeier, I., and Finkelstein, R.R. (2022). ABI5 binding protein2 inhibits ABA responses during germination without ABA-INSENSITIVE5 degradation. Plant Physiol 189, 666–678. 10.1093/plphys/kiac096.

17. Lynch, T.J., Erickson, B.J., Miller, D.R., and Finkelstein, R.R. (2017). ABI5-binding proteins (AFPs) alter transcription of ABA-induced genes via a variety of interactions with chromatin modifiers. Plant Mol Biol 93, 403–418. 10.1007/s11103-016-0569-1.

18. Vittozzi, Y., Krüger, T., Majee, A., Née, G., and Wenkel, S. (2024). ABI5 binding proteins: key players in coordinating plant growth and development. Trends Plant Sci. 10.1016/j.tplants.2024.03.009.

19. Staudt, A.-C., and Wenkel, S. (2011). Regulation of protein function by microProteins. EMBO Rep 12, 35–42.

20. Eguen, T., Straub, D., Graeff, M., and Wenkel, S. (2015). MicroProteins: small size-big impact. Trends Plant Sci 20, 477–482. 10.1016/j.tplants.2015.05.011.

21. Petri, L., Van Humbeeck, A., Niu, H., Ter Waarbeek, C., Edwards, A., Chiurazzi, M.J., Vittozzi, Y., and Wenkel, S. (2025). Exploring the world of small proteins in plant biology and bioengineering. Trends Genet 41, 170–180. 10.1016/j.tig.2024.09.004.

22. Finkelstein, R.R., and Lynch, T.J. (2022). Overexpression of ABI5 Binding Proteins Suppresses Inhibition of Germination Due to Overaccumulation of DELLA Proteins. Int J Mol Sci 23. 10.3390/ijms23105537.

23. Carianopol, C.S., Chan, A.L., Dong, S., Provart, N.J., Lumba, S., and Gazzarrini, S. (2020). An abscisic acid-responsive protein interaction network for sucrose non-fermenting related kinase1 in abiotic stress response. Commun Biol 3, 145. 10.1038/s42003-020-0866-8.

24. Carianopol, C.S., and Gazzarrini, S. (2020). SnRK1α1 Antagonizes Cell Death Induced by Transient Overexpression of Arabidopsis thaliana ABI5 Binding Protein 2 (AFP2). Front Plant Sci 11, 582208. 10.3389/fpls.2020.582208.

25. Martí, E., Gisbert, C., Bishop, G.J., Dixon, M.S., and García-Martínez, J.L. (2006). Genetic and physiological characterization of tomato cv. Micro-Tom. J Exp Bot 57, 2037–2047. 10.1093/jxb/erj154.

26. Altmann, T. (1998). A tale of dwarfs and drugs: brassinosteroids to the rescue. Trends Genet 14, 490–495. 10.1016/s0168-9525(98)01598-4.

27. Roxrud, I., Lid, S.E., Fletcher, J.C., Schmidt, E.D., and Opsahl-Sorteberg, H.G. (2007). GASA4, one of the 14-member Arabidopsis GASA family of small polypeptides, regulates flowering and seed development. Plant Cell Physiol 48, 471–483. 10.1093/pcp/pcm016.

28. Sun, S., Wang, H., Yu, H., Zhong, C., Zhang, X., Peng, J., and Wang, X. (2013). GASA14 regulates leaf expansion and abiotic stress resistance by modulating reactive oxygen species accumulation. J Exp Bot 64, 1637–1647. 10.1093/jxb/ert021.

29. Zimmermann, R., Sakai, H., and Hochholdinger, F. (2010). The Gibberellic Acid Stimulated-Like gene family in maize and its role in lateral root development. Plant Physiol 152, 356–365. 10.1104/pp.109.149054.

30. Zhou, R., Fan, M., Zhao, M., Jiang, X., and Liu, Q. (2022). Overexpression of LtKNOX1 from Lilium tsingtauense in Nicotiana benthamiana affects the development of leaf morphology. Plant Signal Behav 17, 2031783. 10.1080/15592324.2022.2031783.

31. Gellon, G., and McGinnis, W. (1998). Shaping animal body plans in development and evolution by modulation of Hox expression patterns. Bioessays 20, 116–125. 10.1002/(sici)1521-1878(199802)20:2116::Aid-bies4>3.0.Co;2-r.

32. Glaser, T., Jepeal, L., Edwards, J.G., Young, S.R., Favor, J., and Maas, R.L. (1994). PAX6 gene dosage effect in a family with congenital cataracts, aniridia, anophthalmia and central nervous system defects. Nat Genet 7, 463–471. 10.1038/ng0894-463.

33. Fernandez, A.I., Viron, N., Alhagdow, M., Karimi, M., Jones, M., Amsellem, Z., Sicard, A., Czerednik, A., Angenent, G., Grierson, D., et al. (2009). Flexible tools for gene expression and silencing in tomato. Plant Physiol 151, 1729–1740. 10.1104/pp.109.147546.

34. Jin, Y., and Marquardt, S. (2020). Dual sgRNA-based Targeted Deletion of Large Genomic Regions and Isolation of Heritable Cas9-free Mutants in Arabidopsis. Bio Protoc 10, e3796. 10.21769/BioProtoc.3796.

35. Edwards, K., Johnstone, C., and Thompson, C. (1991). A simple and rapid method for the preparation of plant genomic DNA for PCR analysis. Nucleic Acids Res 19, 1349. 10.1093/nar/19.6.1349.

36. Kopylova, E., Noé, L., and Touzet, H. (2012). SortMeRNA: fast and accurate filtering of ribosomal RNAs in metatranscriptomic data. Bioinformatics 28, 3211–3217. 10.1093/bioinformatics/bts611.

37. Bolger, A.M., Lohse, M., and Usadel, B. (2014). Trimmomatic: a flexible trimmer for Illumina sequence data. Bioinformatics 30, 2114–2120. 10.1093/bioinformatics/btu170.

38. Patro, R., Duggal, G., Love, M.I., Irizarry, R.A., and Kingsford, C. (2017). Salmon provides fast and bias-aware quantification of transcript expression. Nat Methods 14, 417–419. 10.1038/nmeth.4197.

39. Dobin, A., and Gingeras, T.R. (2016). Optimizing RNA-Seq Mapping with STAR. Methods Mol Biol 1415, 245–262. 10.1007/978-1-4939-3572-7_13.

40. Love, M.I., Huber, W., and Anders, S. (2014). Moderated estimation of fold change and dispersion for RNA-seq data with DESeq2. Genome Biol 15, 550. 10.1186/s13059-014-0550-8.

41. Santos, A., Colaço, A.R., Nielsen, A.B., Niu, L., Strauss, M., Geyer, P.E., Coscia, F., Albrechtsen, N.J.W., Mundt, F., Jensen, L.J., and Mann, M. (2022). A knowledge graph to interpret clinical proteomics data. Nat Biotechnol 40, 692–702. 10.1038/s41587-021-01145-6.

42. Lazar, C., Gatto, L., Ferro, M., Bruley, C., and Burger, T. (2016). Accounting for the Multiple Natures of Missing Values in Label-Free Quantitative Proteomics Data Sets to Compare Imputation Strategies. J Proteome Res 15, 1116–1125. 10.1021/acs.jproteome.5b00981.

43. Tyanova, S., and Cox, J. (2018). Perseus: A Bioinformatics Platform for Integrative Analysis of Proteomics Data in Cancer Research. Methods Mol Biol 1711, 133–148. 10.1007/978-1-4939-7493-1_7.

44. Šimura, J., Antoniadi, I., Široká, J., Tarkowská, D., Strnad, M., Ljung, K., and Novák, O. (2018). Plant Hormonomics: Multiple Phytohormone Profiling by Targeted Metabolomics. Plant Physiol 177, 476–489. 10.1104/pp.18.00293.

